# Paving the way for plant-based bioproduction of (Z)-13-octadecenoic moth pheromone compounds

**DOI:** 10.64898/2026.05.25.727624

**Authors:** Mengyu Liu, Huijing Li, Huanhuan Jin, Yue Xiao, Panpan He, Chang Tan, Hong-Lei Wang, Jean-Marc Lassance, Christer Löfstedt, Bao-Jian Ding

**Author notes:** Author for correspondence: Baojian Ding, Christer Löfstedt, Mengyu Liu. These authors contributed equally to this work (Mengyu Liu, Huijing Li).

## Abstract

Sex pheromones are eco-friendly agents to control pest insects. Yet, the cost of synthetic pheromone production has restricted their use essentially to high value crops. Plant-based bioproduction of pheromone compounds offers a promising and sustainable alternative, opening up for the deployment of pheromone-mediated pest control also in row crops. The rice leaffolder *Cnaphalocrocis medinalis* uses C18 pheromone compounds and is a major rice pest in South China. We demonstrate that the fatty acid precursor of the main C18 pheromone component of *C. medinalis*, (*Z*)-13-octadecenal, can be biotechnologically produced using the oilseed crop *Camelina sativa*. First, we used deuterium-labelling to establish an elongation-based pathway for pheromone precursor formation from hexadecanoic (palmitic) acid via desaturation to (*Z*)-11-hexadecenoic acid and elongation to (*Z*)-13-octadecenoic acid. Using transient expression in *Nicotiana benthamiana*, we identified a minimal pathway consisting of a Δ11 desaturase (*CmedDES1*), an elongase (*CmedELO1*), and a fatty acyl reductase (*CmedFAR2*) for the heterologous biosynthesis of long-chain pheromone monounsaturated C18 precursors in planta. Co-expression of the insect desaturases *CmedDES1* or *CsupYPAQ* with the Camelina elongase *CsaKCS1* in *Nicotiana benthamiana* resulted in higher production of Z13-18 products compared to co-expression with insect elongases. We finally engineered *Camelina sativa* seeds through seed-specific co-expression of the Z11-16-forming insect desaturase *CsupYPAQ* and *CsaKCS1* elongase, enabling the accumulation of C18 fatty acid pheromone precursors. Our study paves the way for sustainable production of the sex pheromone of *C. medinalis* and other C18 monounsaturated pheromone precursors for crop protection.

## 1. Introduction

Conventional insecticides are widely used to control herbivorous insect populations. However, the rapid development of insecticide resistance in pests has made the pest control increasingly challenging (Liang et al., 2025; Wan et al., 2025; Wolfram et al., 2026). In response to insecticide resistance, farmers tend to increase the doses of insecticide applied per unit area, which leads to public concerns regarding insecticide residues in the food products and adverse effects of pesticides on ecosystems (Köninger et al., 2026).

For lepidopteran pests, one particularly promising strategy to reduce the reliance on conventional insecticides is the use of sex pheromones in IPM to prevent or mitigate damage through approaches like population monitoring, mass trapping or mating disruption (Cork, 2016; Evenden, 2016; Suckling, 2016). Due to the costs associated with pheromone production and application, however, the use of pheromones for mating disruption is limited in row crops like rice. Recently, biological production of pheromones has been put forward as an environmentally friendly and economically attractive option (Löfstedt and Xia, 2021; Petkevicius et al., 2020) which may open up for increased use of pheromone-based population control also in row crops.

Most of the moth pheromone components are volatile fatty acid derivatives that are produced in the female moth’s pheromone gland and released to attract conspecific males (Ando et al., 2004; Tillman et al., 1999). In the insect, the biosynthetic pathway for this so-called Type I moth pheromones starts from *de novo* synthesis of palmitic and stearic acids, followed by desaturation and limited chain-shortening or chain elongation, forming the species-specific carbon chain skeletons. Finally, the modification of the carboxyl group into aldehydes, alcohols, and acetate esters yields the active pheromone components (Jurenka, 2021; Löfstedt et al., 2016).

By reconstitution of a combination of insect-derived genes and genes from plants and yeast it has been possible to produce several moth sex pheromone compounds or their immediate fatty acid precursors in biofactories. For example, (Z)-11-hexadecenoic acid (Z11-16:Acid), the immediate fatty acid precursor for aldehyde, alcohol and acetate pheromone components of the diamond back moth *Plutella xylostella* and many heliothine moths was produced successfully in the oil seed crop *Camelina sativa*. The pheromones derived from plant production showed similar biological activity when compared with chemically produced pheromones in field trapping and mating disruption experiments (Wang et al., 2022). Monitoring and mating disruption with the same compound produced by yeast fermentation yielded equivalent and equally promising results (Betsi et al., 2026; Holkenbrink et al., 2020; Raptopoulos et al., 2025). In addition, the immediate fatty acid precursor of codlemone (E, E)-8,10-dodecadienol was produced in Camelina and showed biological activity comparable with conventionally produced pheromone when tested in field trapping and flight tunnel experiments (Xia et al., 2021).

Rice is the staple food of half of the world’s population (Yuan et al., 2021). In 2023, 800 million tons were produced with China, India and Indonesia being the largest consumers. Rice yield can be severely influenced by insect pests including several species of Lepidoptera among which some of the more important ones belong to the Crambidae family and use aldehyde, acetate, and alcohol C16 and C18 pheromone components (T. Ando and M. Yamamoto, 2025. Internet Database. https://lepipheromone.sakura.ne.jp/lepi_phero_ list_eng.html). The Asiatic rice borer or striped rice borer *Chilo suppresalis* uses (*Z)*-13-octadecenal (Z13-18: Ald) as an important secondary component in addition to the major component (*Z*)-11-hexadecenal (Z11-16 aldehyde) (Beevor et al., 1977; Nesbitt et al., 1975). Another major lepidopteran pest on rice in Asian countries is the rice leaffolder *Cnaphalocrocis medinalis* Guenée (Lepidoptera: Pyralidae). Female *C. medinalis* produce a sex pheromone that consists of (*Z*)-11-octadecenal (Z11-18:Ald) and Z13-18:Ald and in China and Japan (Kawazu et al., 2000; Kawazu et al., 2001; Kawazu et al., 2002; Kawazu et al., 2009) the Z13-18:Ald is the major pheromone component in this species. The larvae can spin silk around the rice leaf longitudinally, leading to folded leaves and they consume the upper epidermis and mesophyll tissues of the leaf, leaving only the lower epidermis, and forming a linear pale white stripe in attacked leaves (Khan et al., 1988). Plants damaged by *C. medinalis* are tempered in their photosynthetic ability, resulting in yield loss (Padmavathi et al., 2013). Mating disruption technology applied in paddy fields has been shown to disrupt the rice leaffolder mating with an average disruption rate of 90%, resulting in a decline in the rice leaffolder population, and an increase in the parasitoid number in the treated fields due to the reduction of chemical insecticides compared with the control fields (Wang et al., 2024).

So far, C18 pheromone compounds with a Z13 double bond have not been biologically produced. To pave the way for biotechnological production of C18 pheromone compounds, in particular these with a Z13 unsaturation, we first elucidated the sex pheromone biosynthetic pathway in the female *C. medinalis* with the aim of identifying enzymes that could be used to reconstitute the biosynthetic pathway in platforms for biotechnological pheromone production. We first identified fatty acids in the pheromone gland that could serve as potential pheromone precursors and performed labeling experiments to study incorporation of deuterated fatty acids into pheromone compounds. After that, by conducting high-throughput sequencing of the female *C. medinalis* pheromone gland and abdominal cuticle transcriptomes, we identified and characterized candidate genes encoding desaturases (DES), long-chain fatty acid elongases (ELO) and fatty acyl reductases (FAR) that are putatively involved in the pheromone biosynthetic pathway of this moth and functionally expressed them using *Nicotiana benthamiana* as a plant expression system. Furthermore, we compared the activity of the rice leaffolder desaturase and elongase to enzymes derived from other sources but with similar activity profiles by expression in the *N. benthamiana* system. Finally, we co-opted *Camelina sativa* as a production platform to demonstrate the feasibility of producing the immediate fatty acid precursor of the (*Z*)-13-octadecenoic pheromone compounds in this oil crop.

## 2. Result

### 2.1 Reconstruction of an elongation-based pathway for C18 pheromone precursor biosynthesis

To establish a framework for pathway reconstruction, we elucidated the biosynthetic origin of the main sex pheromone component in *C. medinalis* using gas chromatography-mass spectrometry (GC-MS). Aside from common saturated and monounsaturated fatty acids, the pheromone gland lipid profile contained several pheromone-related fatty acid precursors, namely (*Z*)-11-hexadecenoic acid (Z11-16:Acid), (*Z*)-11-octadecenoic acid (Z11-18:Acid) and (*Z*)-13-octadecenoic acid (Z13-18:Acid) (Fig. S1). The presence of these intermediates suggested two alternative biosynthetic routes: Δ11 desaturation of C16 acyl precursors followed by chain elongation to C18, or Δ13 desaturation on C18 substrates.

To resolve these possibilities, we performed *in vivo* labeling experiments using deuterium-labeled fatty acid precursors and monitored incorporation of deuterium atoms into pathway intermediates and the final product using GC-MS analysis. Application of D_3_-16:acid resulted in label incorporation in 16:Acid, Z11-16:Acid, Z13-18:Acid and Z13-18:Ald (Fig. 1A). Similarly, deuterium atoms from D_3_-Z11-16:acid were incorporated into Z13-18:Acid and Z13-18:Ald (Fig. 1A). In contrast, no label incorporation from D_3_-18:Acid into Z13-18:Acid or its aldehyde derivative was detected (Fig. 1A). Together, these results support a biosynthetic route in which C16 substrates undergo Δ11 desaturation followed by elongation to C18 (Fig. 1B), establishing an elongation-based pathway for pheromone precursor formation.

**Figure 1.**
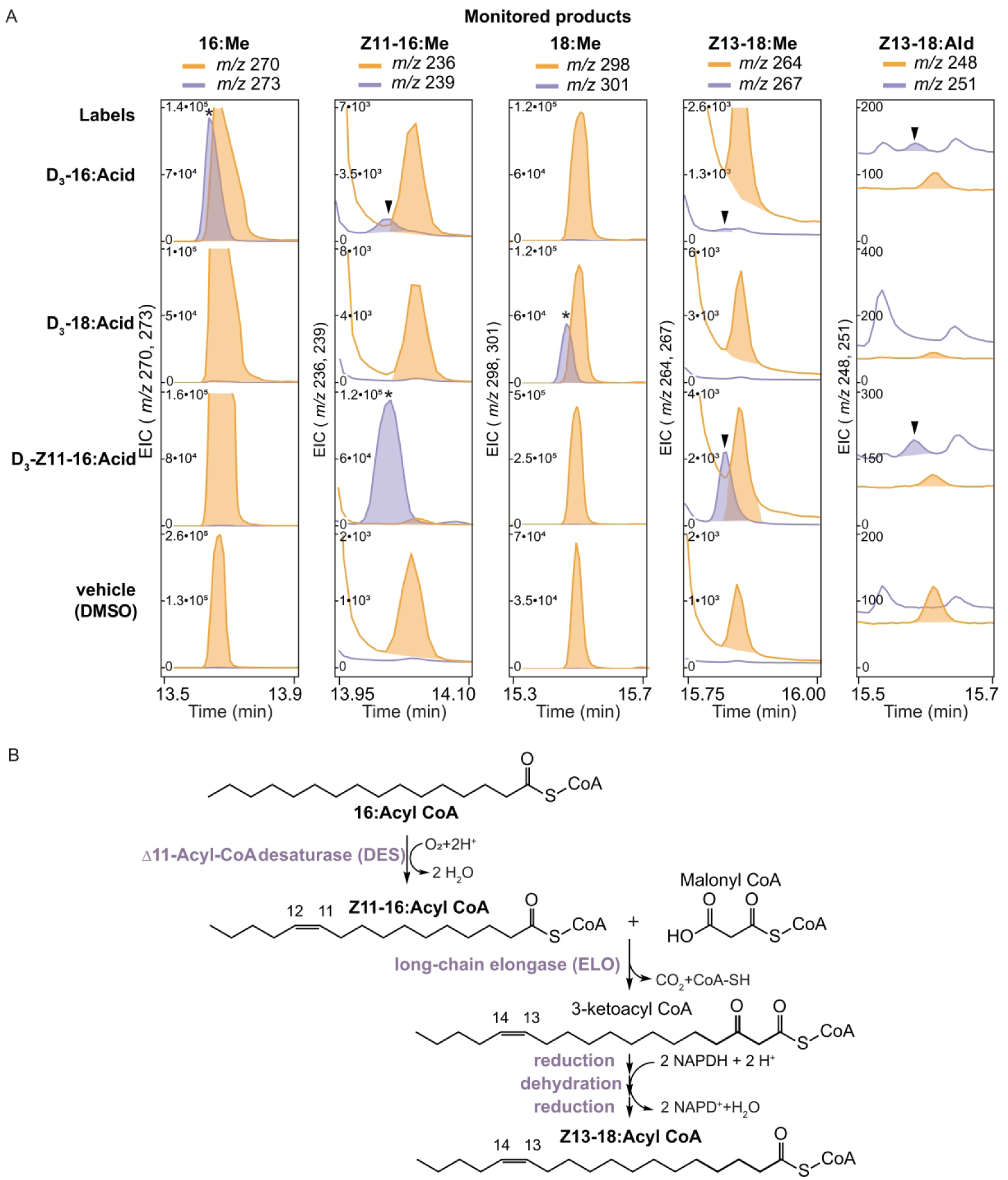
Reconstruction of the biosynthetic pathway for the Δ13-octadecenoic acid pheromone precursor. (A) GC–MS selected ion monitoring (SIM) analysis of deuterium incorporation from labelled fatty acid precursors. Extracted ion chromatograms (EICs) are shown for individual compounds, each displaying the native ion and the corresponding +3 Da deuterium-labelled ion: methyl hexadecanoate (16; *m/z* 270/273), methyl (*Z*)-11-hexadecenoate (Z11-16; *m/z* 236/239), methyl octadecanoate (18; *m/z* 298/301), methyl (Z)-13-octadecenoate (Z13-18; *m/z* 264/267), and (*Z*)-13-octadecenal (Z13-18; m/z 248/251). Treatments included [16,16,16-²H₃] hexadecanoic acid (D₃-16), [18,18,18-²H₃] octadecanoic acid (D₃-18), [16,16,16-²H₃] (*Z*)-11-hexadecenoic acid (D₃-Z11-16) –all denoted by an asterisk – and vehicle control (DMSO). Incorporation of deuterium is indicated by the appearance of +3 Da ions (arrowheads). For each compound, only the selected native and corresponding labelled ions are displayed. Native compounds are shown as orange traces and deuterium-labelled compounds as purple traces. (B) Proposed biosynthetic pathway towards Z13-18:Acyl precursor in *C. medinalis*. Hexadecanoyl-CoA is converted to (*Z*)-11-hexadecenoyl-CoA by a Δ11 desaturase (DES), followed by elongation to (*Z*)-13-octadecenoyl-CoA via a fatty acid elongase (ELO) system involving condensation with malonyl-CoA (rate-limiting step), reduction, dehydration and reduction steps. In the gland, the acyl intermediate is subsequently converted to the corresponding alcohol and aldehyde by reduction and oxidation (steps not depicted).

### 2.2 Functional reconstitution of desaturation and elongation modules in Nicotiana benthamiana

To identify candidate enzymes for pathway reconstruction toward the production of Z13-18:Ald, we performed RNA sequencing (RNA-Seq) of female pheromone gland and abdominal cuticle tissues, which differ in their ability to biosynthesize the octadecenyl substrate (Fig. 2A). Differential expression analysis revealed clear separation between the two tissues (Fig. 2B), where biological replicates clustered tightly by tissue type, indicating distinct transcriptional programs. In total, 1,572 differentially expressed genes (DEGs) were identified in the pheromone gland relative to the abdominal cuticle, including 1,072 upregulated genes (FDR < 0.05). KEGG enrichment analysis further showed that these DEGs were significantly enriched in pathways associated with fatty acid biosynthesis, biosynthesis of unsaturated fatty acids, and fatty acid metabolism (Fig. 2C).

**Figure 2.**
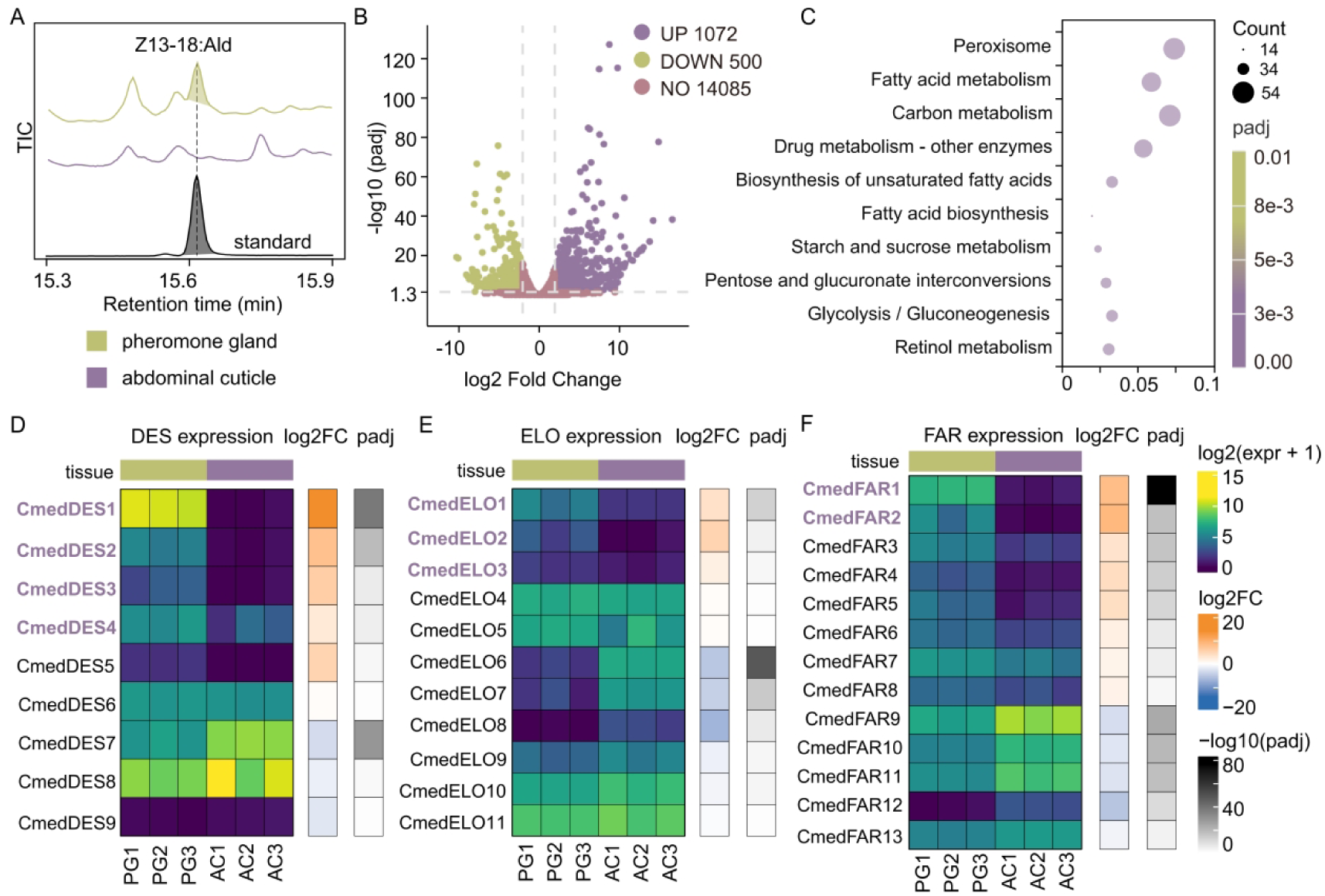
Tissue-specific pheromone production and transcriptomic analysis of pheromone gland in *C. medinalis*. (A) GC–MS analysis of (*Z*)-13-octadecenal (Z13-18) in female tissues. Total ion chromatograms (TICs) are shown for pheromone gland (PG; top), abdominal cuticle (AC; middle), and Z13-18 standard (bottom). (B) Volcano plot showing differentially expressed genes (DEGs) between PG and AC. The x-axis represents log_2_ fold change and the y-axis shows the -log_10_ (adjusted P-value). Genes significantly upregulated in PG are shown in purple, and downregulated genes in light green (FDR ≤ 0.05 and |log_2_FC|≥ 1). (C) Kyoto Encyclopedia of Genes and Genomes (KEGG) pathway analysis of genes upregulated in PG relative to AC. Only significantly enriched pathways (adjusted P < 0.05) are shown. Dot size indicates gene count and color represents adjusted P value. (D-F) Heatmaps showing expression profiles of fatty acid desaturase (DES) (D), elongase chain elongase (ELO) (E), and fatty acyl reductase (FAR) (F) gene families in PG and AC. Expression values are shown as log_2_(FPKM +1). Genes highlighted in red indicate candidates selected for functional characterization.

Based on the predicted biosynthesis pathway (Fig.1B), we searched for candidate genes encoding fatty acid desaturase (DES) and long-chain fatty acid elongase (ELO) enriched in the pheromone gland (Fig. 2D, E). We prioritized candidates according to their expression levels and similarity to functionally characterized enzymes involved in fatty acid modification. Four DES candidates (*CmedDES1* to *CmedDES4*) were identified as highly expressed in the pheromone gland (Fig. 2D). Quantitative RT–PCR confirmed their enrichment relative to other female tissues (Fig. S2). Phylogenetic analysis indicated that *CmedDES1* clusters within a Lepidoptera-specific Δ11 desaturase clade and is closely related to genes encoding enzymes known to convert C16 substrates into Z11-16 intermediates, whereas the remaining DES candidates grouped within clades associated with alternative or uncharacterized desaturation activities (Fig. S3).

To assess their functional roles, candidate desaturases were expressed in a *Nicotiana benthamiana* (*N. benthamiana*) transient expression system. Expression of *CmedDES1* resulted in the production of Z11-16-derived products, consistent with Δ11 desaturation activity on C16 substrates (Fig. 3A, C). In addition, *CmedDES1* also conferred the ability to catalyze the production of Z11-18:Acid from C18 substrates, albeit at lower levels (Fig. 3A, D). In contrast, the other DES candidates showed either different from the sought activity (e.g., Δ9) or negligible activity toward C16 and C18 substrates (Fig. 3). These results identify CmedDES1 as the primary desaturase responsible for generating the Z11-16 intermediate required for downstream elongation.

**Figure 3.**
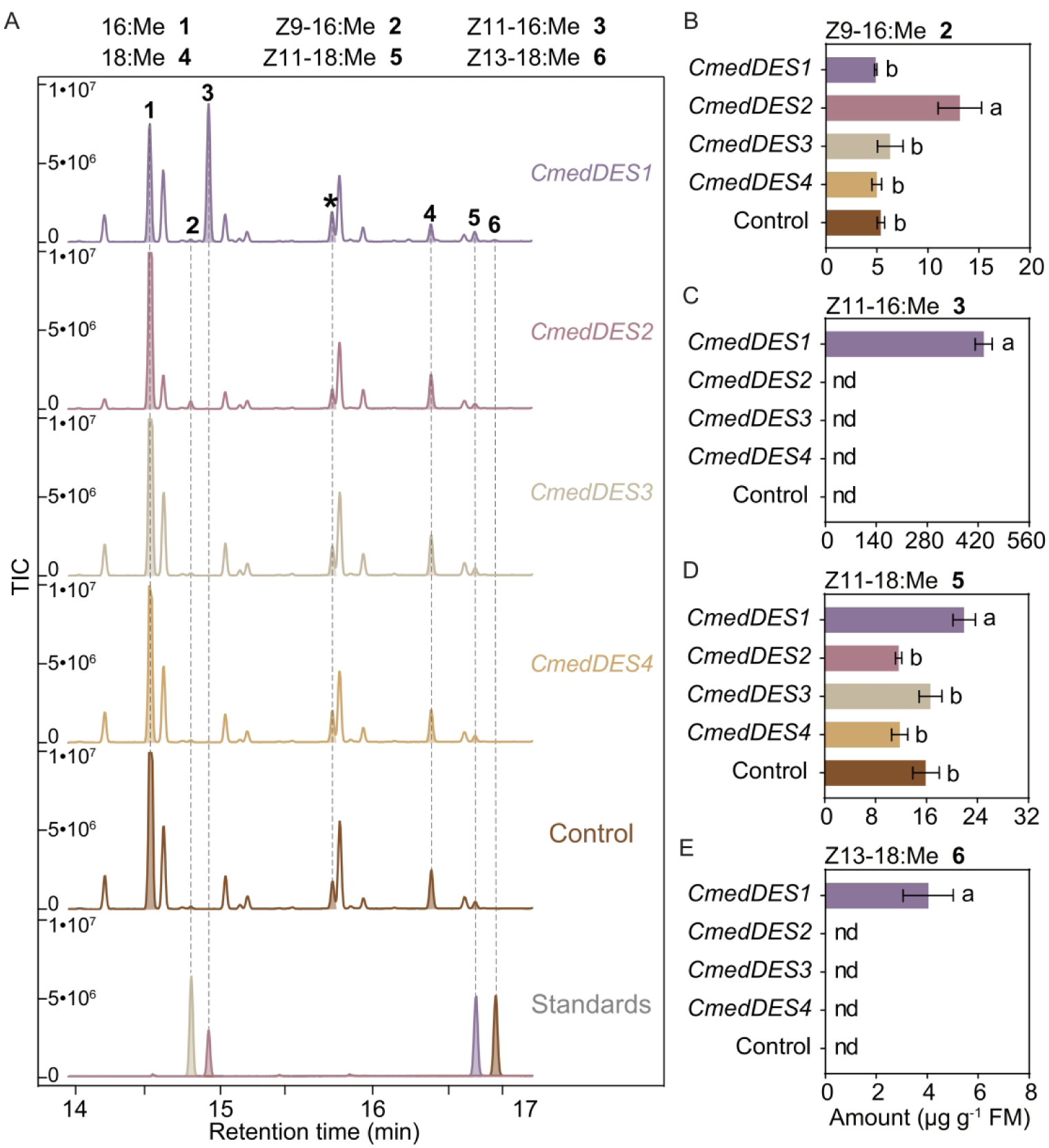
Functional characterization of desaturase candidates from *C. medinalis*. (A) GC–MS analysis of fatty acid methyl esters (FAMEs) produced in *Nicotiana benthamiana* leaves expressing *C. medinalis* desaturases (CmedDES1-4). Total ion chromatograms (TICs) are shown for each construct and an empty vector control. Peaks are annotated as follows: 16:Me (1), Z9-16:Me (2), Z11-16:Me (3), 18:Me (4), Z11-18:Me (5), and Z13-18:Me (6). The asterisk indicates the internal standard (Z10-17:Me, 10 µg spiked in). The bottom trace shows TICs of authentic standards. (B-E) Quantification of FAME products in *N. benthamiana* leaves expressing the indicated desaturases. Mean levels (±SE, n = 5) of Z9-16:Me (B), Z11-16:Me (C), Z11-18:Me (D), Z13-18:Me (E) are shown. Metabolite level were measured 72 h after agroinfiltration. nd, not detected. Different letters indicate statistically significant differences among different treatments (one-way ANOVA followed by Tukey’s multiple comparison test; P < 0.05).

Notably, trace amounts of Z13-18:Me were detected upon expression of *CmedDES1* alone, demonstrating that endogenous elongase activity in *N. benthamiana* can extend Z11-16 intermediates to C18:1(Fig. 3A, E). However, the low accumulation of Z13-18 products indicates that elongation is a limiting step, motivating the introduction of dedicated elongase modules to enhance pathway flux. To address this limitation, we next evaluated enzymatic activity in planta by transiently co-expressing candidate *ELO* genes from *C. medinalis* with *CmedDES1* in *N. benthamiana* leaves via agroinfiltration and screened for the production of Z13-18 products (Fig. 4A). The combination of *CmedDES1* and *CmedELO1* resulted in a marked increase in the production of Z13-18 products compared to expression of *CmedDES1* alone, confirming its role in extending Z11-16 intermediates to C18:1 (Fig. 4D). Other *CmedELO* candidates showed lower or negligible activity under the same conditions (Fig. 4). Together, these results demonstrate that CmedDES1 and CmedELO1 constitute a functional desaturation–elongation module capable of producing C18 unsaturated fatty acids in planta, establishing a blueprint for reconstructing the metabolic pathway to produce Z13-18 fatty acid derivatives.

**Figure 4.**
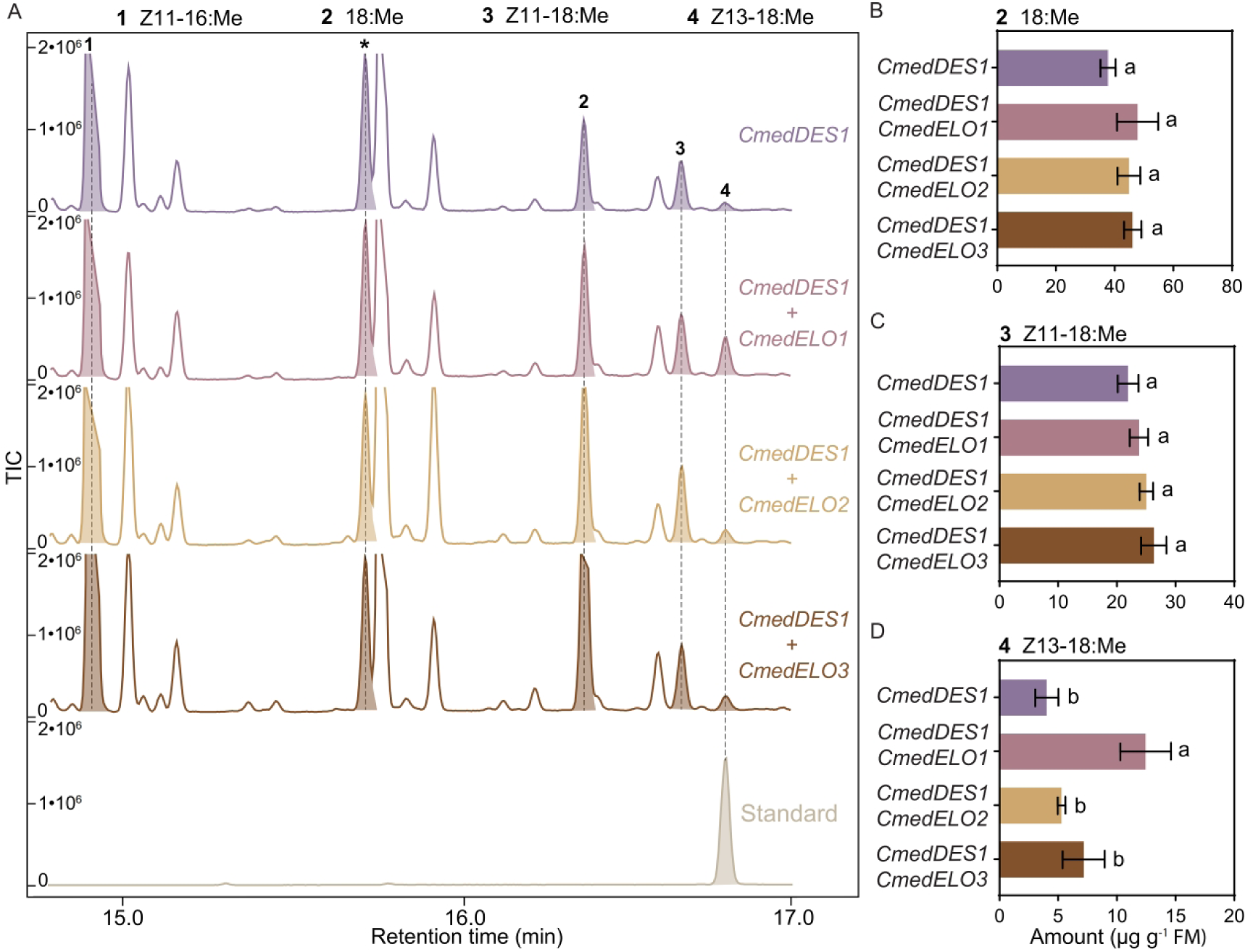
Functional characterization of elongases from *C. medinalis*. (A) GC–MS analysis of fatty acid methyl ester (FAME) profiles from *N. benthamiana* leaves expressing *CmedDES1* alone or in combination with *C. medinalis* elongases (CmedELO1-3). Total ion chromatograms (TICs) are shown for each construct. Peaks are annotated as follows: Z11-16:Me (1), 18:Me (2), Z11-18:Me (3), and Z13-18:Me (4). The asterisk indicates the internal standard (Z10-17:Me, 10 μg spiked in). The bottom trace shows the TIC of an authentic Z13-18:Me standard. (B-D) Quantification of FAME products in *N. benthamiana* leaves expressing the indicated constructs. Mean levels (±SE, n = 5) of 18:Me (B), Z11-18:Me (C), Z13-18:Me (D) are shown. Metabolite levels were measured 72 h after agroinfiltration. nd, not detected. Different letters indicate statistically significant differences among treatments (one-way ANOVA followed by Tukey’s multiple comparison test; P < 0.05).

### 2.3 Validation of a C18-compatible reductase module enables downstream pathway completion

To complete the reconstructed pathway, we next searched for fatty acyl-coA reductases (FARs) in *C. medinalis*. Two candidates (*CmedFAR1* and *CmedFAR2*) were highly expressed in the pheromone gland (Fig. 2F). Phylogenetic analysis placed *CmedFAR2* within the Lepidoptera-specific pgFAR clade, which includes several functionally characterized enzymes implicated in the production of long-chain fatty alcohol pheromone precursors (Fig. 5A). To assess their functional role, each FAR candidate was then transiently co-expressed in *N. benthamiana* leaves together with *CmedDES1* and *CmedELO1* (Fig. 5B-F). Co-expression of *CmedFAR2* resulted in the production of saturated and monounsaturated primary alcohols including (*Z*)-13-octodecenol (Z13-18:OH), demonstrating its ability to catalyze the reduction of both C16 and C18 fatty acyl substrates (Fig. 5F). In contrast, CmedFAR1 showed no activity toward these substrates under the same conditions (Fig. 5E).

**Figure 5.**
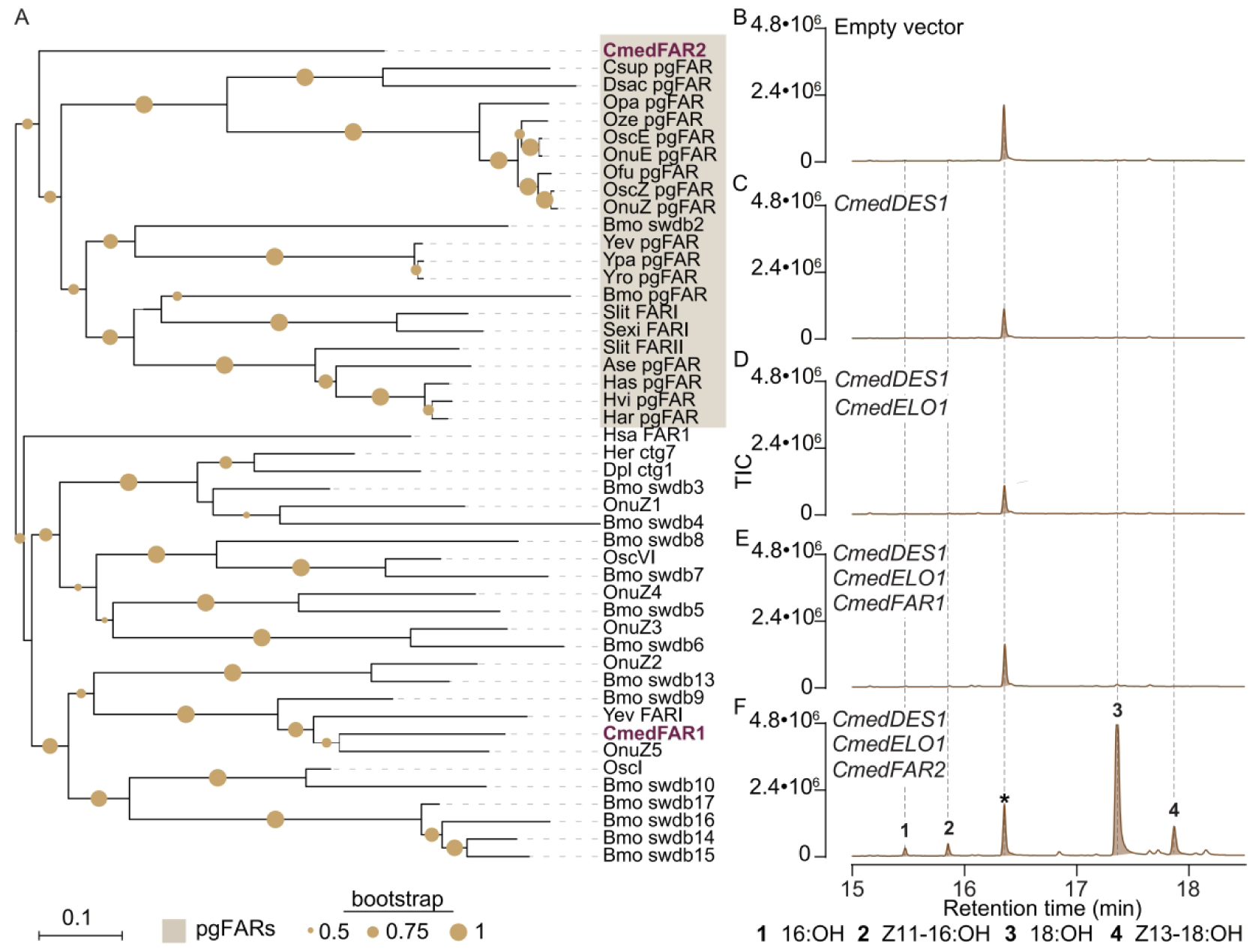
Phylogenetic analysis and functional characterization of candidate fatty acyl reductases (FARs) from *C. medinalis.* (A) Maximum-likelihood phylogenetic tree of candidate *C. medinalis* FARs (CmedFARs) and homologous proteins from other insect species. The clade designated “pgFARs” represents a Lepidoptera-specific group containing FARs involved in moth pheromone biosynthesis. Bootstrap support are indicated at branch nodes (B) GC-MS analysis of fatty alcohol production in *N. benthamiana* leaves expressing combinations of candidate pathway genes. Total ion chromatograms (TICs) are shown for empty vector (control), CmedDES1 alone, CmedDES1 + CmedELO1, and co-expression with candidate FARs (CmedFAR1 or CmedFAR2). Peaks are annotated as follows: 16:OH (1), Z11-16:OH (2), 18:OH (3), and Z13-18:OH (4). Co-expression of CmedDES1 and CmedELO1 with CmedFAR1 results in the production of Z13-18:OH, consistent with reduction of the corresponding acyl precursor. The asterisk indicates the internal standard (17:OH, 10 nmol spiked in).

These results confirm that CmedFAR2 functions as a C18-compatible reductase capable of converting elongated fatty acid intermediates into corresponding alcohols, thereby enabling downstream pathway completion. Together with the desaturation–elongation module, this establishes a functional biosynthetic framework for the heterologous biosynthesis of long-chain pheromone alcohol precursors in planta.

### 2.4 Plant-derived elongase enhances production of non-native C18 unsaturated fatty acids

To further optimize the elongation step, we evaluated whether endogenous plant elongases could improve the conversion of Z11-16 intermediates to C18 products. To ensure robust substrate supply, we employed two Δ11 desaturases, CmedDES1 (*this study*) and CsupYPAQ (Xia et al., 2022), both of which catalyze the formation of Z11-16 intermediates from C16 substrates in planta. These enzymes were used interchangeably as upstream modules to assess elongase performance independently of desaturation efficiency.

In plants, the fatty acid elongation cycle is catalyzed by 3-keto acyl-CoA synthases (KCSs), which determine substrate specificity in the elongation cycle. As our goal was to reconstruct the biosynthetic pathway in the oilseed crop *Camelina sativa* (*C. sativa*), we screened the *Camelina* genome for homologs of *Arabidopsis thaliana* (*A. thaliana*) genes responsible for the condensation activity of elongases involved in producing C16-C20 fatty acids, namely *FAE1* and *KCS1* (Brenda *et al*. 2006). Phylogenetic analysis identified several candidate *KCS* genes, among which *Csa03g002110, Csa17g001600*, and *Csa10g007610* were most similar to *AtFAE1* and *AtKCS1* (Fig. S4). In this manuscript, *Csa03g002110, Csa17g001600*, and *Csa10g007610* will be referred to as *CsaKCS1, CsaKCS1like* and *CsaFAE1,* respectively.

To compare elongase performance, candidate plant KCS genes were co-expressed with desaturase modules in *Nicotiana benthamiana* and assessed for their ability to produce Z13-18 products (Fig. S5A). Among the five plant KCSs tested, expression of *CsaKCS1* resulted in the highest accumulation of Z13-18 products (Fig. S5D). Next, we performed pairwise combinations of desaturase and elongase modules by co-expressing *CmedDES1* and *CsupYPAQ* with *CmedELO1* or *CsaKCS1* (Fig. 6A). No significant difference in Z11-16:Me production was observed between *CmedDES1*/*CmedELO1* and *CsupYPAQ*/*CmedELO1* combinations, indicating comparable catalytic activity between the two desaturases in planta (Fig. 6A, B). However, the highest titer of Z13-18-derived products was obtained with the combination of *CsupYPAQ* with *CsaKCS1* (Fig. 6A, E). Notably, co-expression of the *Camelina* elongase CsaKCS1 resulted in significantly higher accumulation of Z13-18 products compared to co-expression with the insect elongase CmedELO1 (Fig. 6E). This increase was accompanied by a corresponding reduction in Z11-16 intermediates,when *CsaKCS1*, indicating enhanced flux through the elongation step (Fig. 6B).

**Figure 6.**
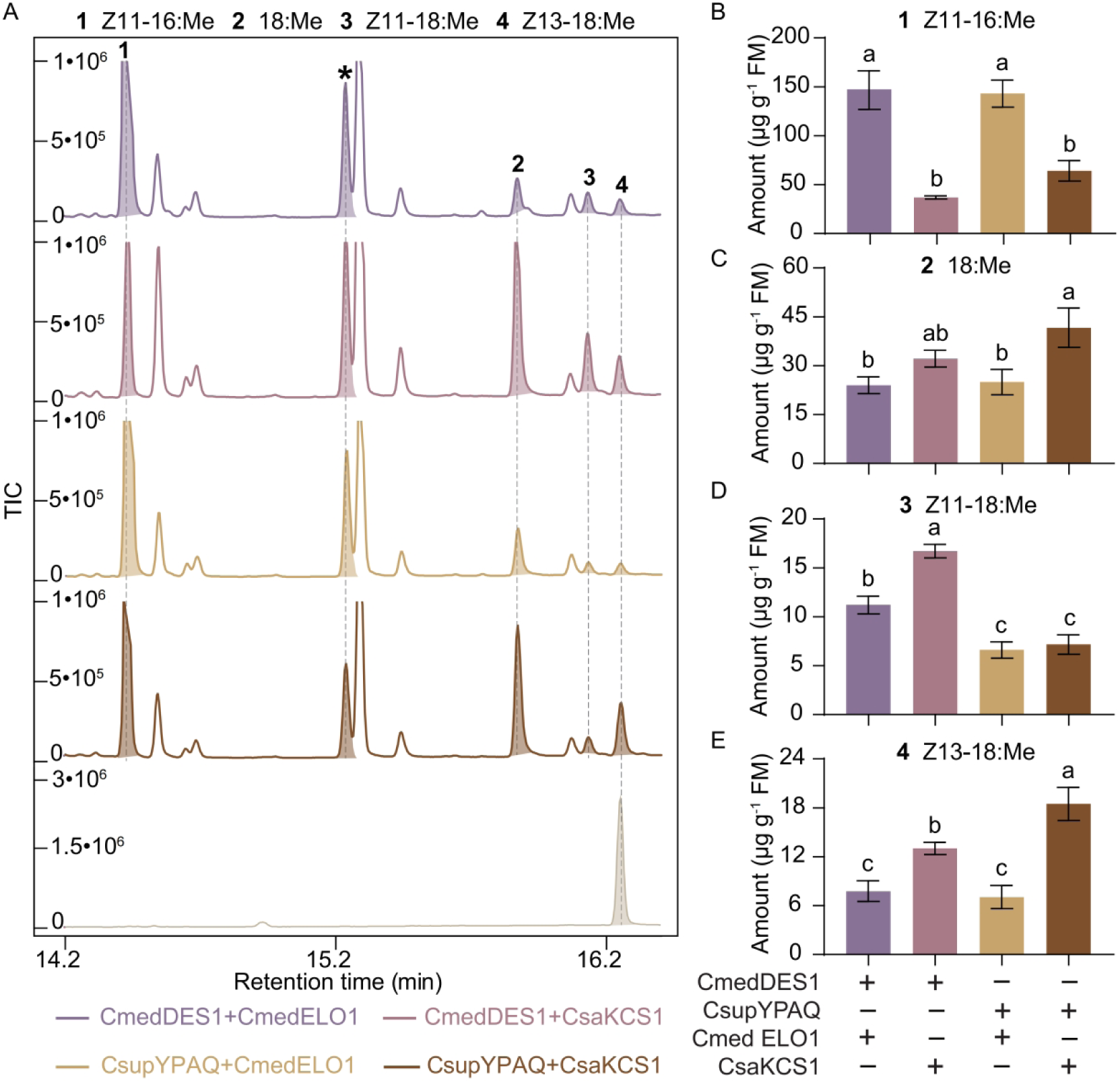
Identification of optimal desaturase-elongase modules for pathway reconstruction. (A) GC–MS analysis of fatty acid methyl ester (FAME) profiles from *N. benthamiana* leaves co-expressing desaturases (CmedDES1 or CsupYPAQ) with elongases (CmedELO1 or CsaKCS1). Total ion chromatograms (TICs) are shown for each combination. Peaks are annotated as follows: Z11-16:Me (1), 18:Me (2), Z11-18:Me (3), and Z13-18:Me (4). The asterisk indicates the internal standard (Z10-17:Me, 10 μg spiked in). The bottom trace shows the TIC of an authentic Z13-18 standard. (B–E) Quantification of FAME products in *N. benthamiana* leaves expressing the indicated constructs. Mean levels (± SE, n = 4) of Z11-16:Me (B), 18:Me (C), Z11-18:Me (D), and Z13-18:Me (E) are shown. Metabolite levels were measured 72 h after agroinfiltration. Different letters indicate statistically significant differences among treatments (one-way ANOVA followed by Tukey’s multiple comparison test; P < 0.05).

Together, these results demonstrate that plant-derived elongases can more efficiently process non-native unsaturated substrates than their insect counterparts in a plant cellular environment. This highlights the importance of host–enzyme compatibility in metabolic pathway reconstruction and suggests that leveraging endogenous or host-adapted enzymes can improve the efficiency of engineered lipid pathways.

### 2.5 Seed-specific metabolic rewiring enables stable accumulation of pheromone precursors in *C. sativa*

To establish a scalable production platform, we next implemented the reconstructed pathway in the oilseed crop *C. sativa* (Fig.7). To enhance the availability of C16 substrates for downstream modification, we first redirected carbon flux toward the cytosolic acyl-CoA pool by expressing a plastidial thioesterase (*FatB1*), thereby increasing the supply of C16 fatty acid precursors. To enable pheromone precursor biosynthesis in seeds, we introduced desaturation and elongation modules under the control of a seed-specific *napin* gene promoter. Transgenic Camelina lines expressing *CsupYPAQ* either with the elongase *CsaKCS1* (Csa_Z13-18) or alone (Csa_Z11-16) were generated and advanced to the T2 generation (Fig. 7A). Importantly, these transgenic lines exhibited normal growth and seed development, with no detectable penalties in plant morphology or seed yield under greenhouse conditions (Fig. 7A; Fig. S6).

**Figure 7.**
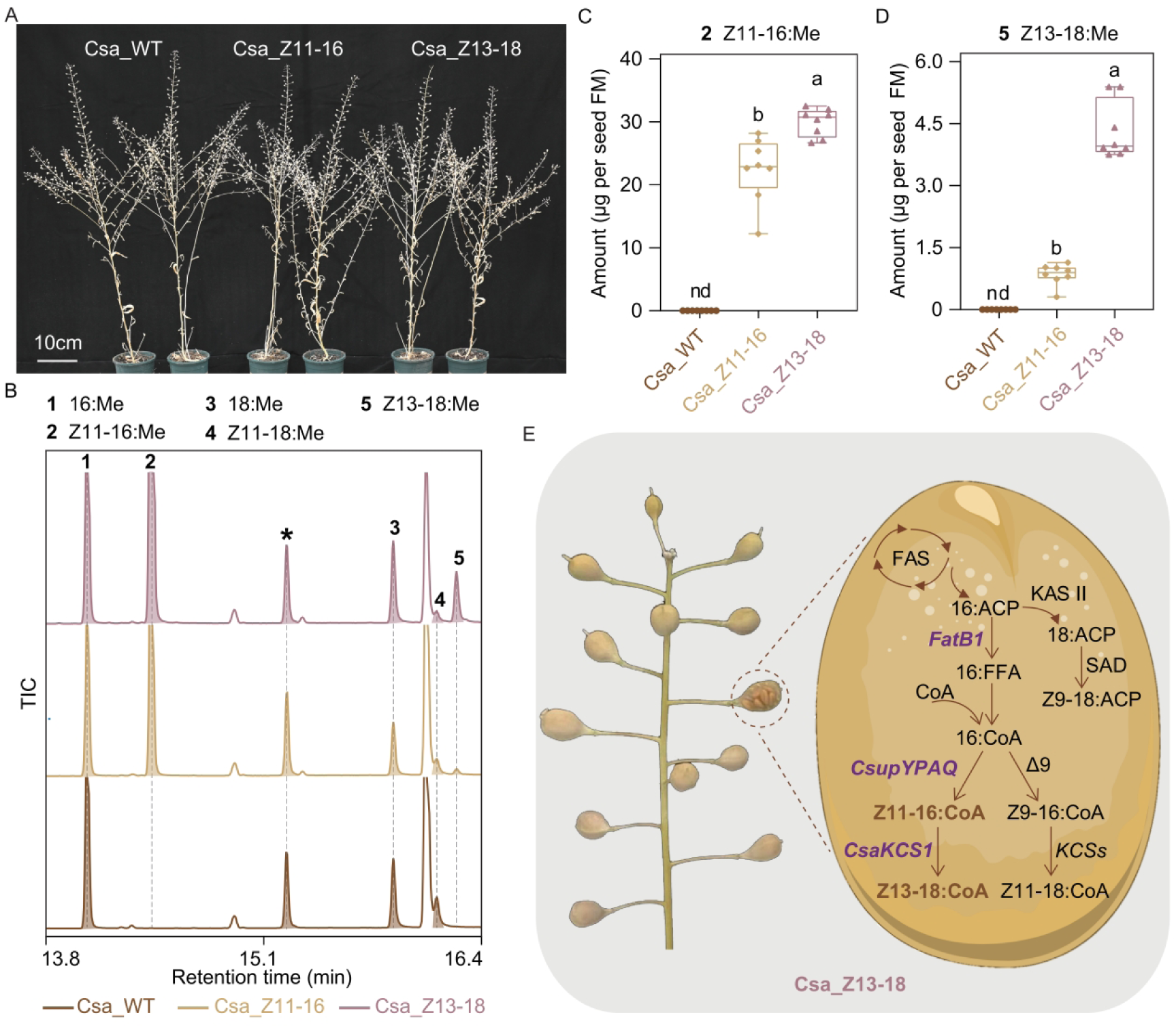
Metabolic engineering of Z13-18 pheromone precursor biosynthesis in *Camelina sativa*. (A) Phenotypes of transgenic *Camelina sativa* lines engineered to produce Z11-16 or Z13-18 fatty acid derivatives, compared with wild-type (WT) plants. Images were taken from 80-day-old plants. Scale bar = 10 cm. (B) GC–MS analysis of fatty acid methyl ester (FAME) profiles from seeds of WT, Csa_Z11-16, and Csa_Z13-18 lines. Total ion chromatograms (TICs) are shown. Peaks are annotated as follows: 16:Me (1), Z11-16:Me (2), 18:Me (3), Z11-18:Me (4), and Z13-18:Me (5). The asterisk indicates the internal standard (Z10-17:Me, 10 μg spiked in).(C, D) Quantification of FAME products in *Camelina* seeds. Mean levels (± SE, n = 8) of Z11-16 (C) and Z13-18 (D) are shown. nd, not detected. Different letters indicate statistically significant differences among lines (one-way ANOVA followed by Tukey’s multiple comparison test; P < 0.05). (E) Schematic representation of the engineered pathway for production of Z13-18 fatty acid derivatives in *Camelina* seeds. Introduced enzymes are indicated in purple, and endogenous enzymes are shown in black. ACP, acyl carrier protein; FAS, fatty acid synthase; FFA, free fatty acid; CoA, coenzyme A; KCS, β-ketoacyl-CoA synthase; SAD, stearoyl-ACP desaturase.

We used GC-MS analysis to examine the fatty acid profile of mature seeds and monitored the accumulation of non-native unsaturated fatty acids, including (Z)-11-hexadecenoic acid (Z11-16:Acid) and (Z)-13-octadecenoic acid (Z13-18:Acid) (Fig. 7B). In both lines, the expression of *CsupYPAQ* significantly increased the content of Z11-16:Acid (Fig. 7C). Furthermore, the line expressing the plant-derived elongase *CsaKCS1* showed significantly higher levels of Z13-18 products compared to the transgenic line only expressing the constitutive elongation complex (Csa_Z11-16) (Fig. 7D), consistent with observations from transient expression assays in *N. benthamiana*. Accumulation levels reached up to 5.5 µg per seed, demonstrating efficient flux through the engineered pathway (Fig. 7D).

Taken together, these results demonstrate that seed lipid metabolism can be effectively rewired to produce non-native C18 unsaturated fatty acids and establish *Camelina sativa* as a viable platform for the stable production of C18 monounsaturated pheromone precursors (Fig. 7E).

## 3. Discussion

In this study, we elucidated the biosynthetic pathway underlying production of the major sex pheromone component of *Cnaphalococris medinalis* and reconstructed this pathway in plant systems for biotechnological production of C18 pheromone precursors. Using precursor analysis and stable-isotope labeling, we demonstrated that pheromone biosynthesis proceeds through Δ11 desaturation of C16 fatty acids followed by chain elongation to generate the Z13-18 intermediate. Functional characterization of desaturation, elongation, and reduction modules in *Nicotiana benthamiana* enabled reconstruction of the core biosynthetic pathway *in planta*. Furthermore, stable expression of optimized pathway elements was achieved to demonstrate the production of non-native C18 unsaturated fatty acid precursors, establishing an oilseed-based of the major *C. medinalis* pheromone component using *Camelina sativa* as a plant biofactory. Together, these findings expand the repertoire of biologically producible moth pheromone compounds and providing a framework for scalable and sustainable pheromone-based pest control in row crops.

The *C. medinalis* sex pheromone was identified already in the 1990’s (Rao et al., 1995), but the biosynthetic pathway leading to its production had yet to be elucidated. Here, we established via precursor analysis and deuterium-labeling that biosynthesis of Z13-18:Ald proceeds through Δ11 desaturation of palmitic acid, followed by chain elongation to C18 intermediates, and final conversion of the acid into the volatile Z13-18:Ald pheromone component. This pathway architecture could not be inferred a priori because moth pheromone biosynthesis frequently involves highly specialized and evolutionarily diversified fatty acyl desaturases. Lepidopteran genomes contain large repertoires of desaturase genes, many of which encode unusual regioselectivities and substrate specificities, including activities rarely observed in other organisms (e.g., Δ5, Δ6, Δ14, Δ10) (Lassance et al., 2021). Consequently, both Δ11 desaturation followed by elongation and direct Δ13 desaturation of C18 substrates represented plausible biosynthetic scenarios for Z13-18 production. Expression analyses and functional characterization identified CmedDES1 as the key enzyme generating the Z11-16 intermediate required for formation of C18 pheromone precursors. Phylogenetically, this desaturase clusters within the Lepidoptera-specific Δ11 clade (Lienard et al., 2008), a desaturase lineage containing enzymes which are involved in the biosynthesis of moth pheromone components (Löfstedt et al., 2016). However, because the FAD family in moths has experienced substantial expansion and neofunctionalization, the resulting broad functional diversity highlights the importance of direct biochemical validation, as sequence similarity-based classification alone is insufficient to predict catalytic activity or involvement in pheromone biosynthesis. Together, our findings establish an elongation-based route for C18 pheromone precursor formation and define the minimal desaturation module required for heterologous pathway reconstruction in plants.

Elongases are widely present in various organisms, including animals, plants and microorganisms, primarily regulating the chain length of fatty acids (Guillou et al., 2010). The fatty acid elongation reaction comprises four steps: condensation, first reduction, dehydration, and second reduction. In particular, ELO is the rate-limiting condensation enzyme acting in the first step (Leonard et al., 2004). Compared to moth desaturases and reductases, relatively little is known about elongases involved in insect pheromone biosynthesis. Notable exceptions are Elo68a and Jamesbond. Elo68a, which is specifically expressed in the reproductive organs of male *Drosophila melanogaster* fruitflies, was the first ELO gene identified in insects. Biochemically, Elo68a extends myristoleic and palmitoleic acids, and mutations affecting it reduce the content of *cis*-vaccenyl acetate in males (Chertemps et al., 2005). Jamesbond (also known as bond) which encodes a very-long-chain fatty acid elongase, plays an essential role in the biosynthesis of the male sex pheromone (3*R*,11*Z*,19*Z*)-3-acetoxy-11,19-octacosadien-1-ol (CH503) (Ng et al., 2015). So, with only a limited number of insect elongases functionally characterized, prediction of substrate specificity from sequence remains difficult. This is particularly relevant for pathways involving C18 intermediates, such as those of *C. medinalis*. Our results identified CmedELO1 as an elongase capable of extending Z11-16 to Z13-18, thereby enabling formation of the Z13-18 precursor required for pheromone biosynthesis.

The detection of trace Z13-18 products following expression of CmedDES1 alone further suggested that elongation represented a limiting step during pathway reconstruction *in planta*. Consistent with this interpretation, co-expression of dedicated elongase modules substantially increased accumulation of Z13-18 derivatives. Noteworthy, there are two distinct families of fatty acid elongase condensing enzymes involved in the first step of chain elongation. In mammals, yeast, protists and other eukaryotes including insects, the condensation step is catalyzed by ELO elongases, whereas plants and some protists primarily use unrelated KCSs that are typified by Arabidopsis fatty acid elongase 1 (FAE1). This can explain our finding that the insect elongase CmedELO1 works but was not be the favored choice for biosynthesis in planta. Notably, the Camelina elongase CsaKCS1 consistently outperformed the insect elongase CmedELO1 in the *Nicotiana benthamiana* expression system, resulting in significantly higher accumulation of Z13-18 products. This increase was accompanied by a corresponding reduction in Z11-16 intermediates, indicating enhanced flux through the elongation step and suggesting that the elongation step is a bottle neck in the production. This finding highlights the importance of host–enzyme compatibility in heterologous pathway engineering and suggests that endogenous or host-adapted enzymes may provide superior metabolic performance compared to phylogenetically unrelated enzymes. More broadly, our results demonstrate that efficient reconstruction of specialized lipid pathways may depend not only on identifying biosynthetic enzymes that carry relevant activity in the native organism, but also on optimizing compatibility with host metabolic networks.

Comparison of CmedDES1 with the previously characterized Δ11 desaturase CsupYPAQ further demonstrated that efficient reconstruction of pheromone pathways may require combining biosynthetic modules from different species. Although both insect enzymes catalyzed formation of Z11-16 intermediates, co-expression of CsupYPAQ with the Camelina elongase CsaKCS1 consistently resulted in higher accumulation of Z13-18 products in planta. Previous studies have similarly shown that moth Δ11 desaturases can differ substantially in catalytic efficiency and substrate utilization despite close phylogenetic relationships. These findings highlight the modular nature of pheromone biosynthetic pathways and illustrate how synthetic biology approaches can be used to optimize pathway performance through combinatorial assembly of enzymes from different organisms. Rather than reconstructing native pathways enzyme-by-enzyme, heterologous production platforms may benefit from selecting biosynthetic modules based on catalytic compatibility and metabolic performance within the host environment.

The rice leaffolder *C. medinalis* is a major pest of rice cultivation throughout Asia. The infestation occurs throughout all growth stages of rice, affecting plant growth and fertility, with the most significant impact observed during the reproductive stage (Padmavathi et al., 2013). In the past decades, control of *C. medinalis* has relied mainly on pesticides. However, repeated insecticide applications can produce undesirable ecological consequences, including mortality of natural enemies of *C. medinalis* and non-target insects (Gurr et al., 2012). In addition, the selective pressure generated by insecticide treatments is known to lead to resistance, including in *C. medinalis* (Li et al., 2026; Subhagan et al., 2026; Subhagan et al., 2025). As an alternative to conventional pesticides, pheromone-mediated mating disruption offers an environmentally friendly alternative. Mating disruption applied in paddy fields has been shown to disrupt the rice leaffolder mating with an average disruption rate of 90%, resulting in a substantial decline in rice leaffolder populations, while decreasing insecticide use and therefore promoting natural enemy abundance, with greater numbers of parasitoids in treated fields compared with control fields (Wang *et al*. 2024; Yang et al., 2025). However, the high cost of chemically synthesized pheromones has limited large-scale deployment of pheromone-based pest control in row crops such as rice.

Our results demonstrate that *Camelina sativa* can serve as a stable oilseed platform for production of C18 pheromone precursors. Given typical Camelina seed and oil yields, the production levels obtained here suggest that biologically produced pheromone precursors could support pheromone deployment across substantial cultivation areas. Given that mating disruption to control rice leaffolder requires 120 g of active ingredient, we calculated that growing of one hectare of engineered Camelina will produce enough precursors for mating disruption formulation to control 40 ha of rice field. In addition, the simultaneous accumulation of multiple pheromone precursor molecules within the same seed oil may simplify downstream processing and broaden applicability across geographically distinct pheromone blends. More broadly, these findings support the feasibility of plant-based biomanufacturing for sustainable crop protection and highlight the potential of oilseed metabolic engineering as a scalable alternative to petrochemical pheromone synthesis.

In addition to reconstruction of desaturation and elongation modules, downstream conversion of elongated fatty acid intermediates was enabled by the CmedFAR2. Unlike many previously characterized moth pheromone gland reductases, which preferentially reduce C14 or C16 unsaturated substrates, CmedFAR2 showed strong activity toward C18 monounsaturated fatty acids, including Z13-18:Acid. This expands the repertoire of characterized pheromone biosynthetic reductases and provides a suitable enzymatic module for biosynthesis of long-chain unsaturated alcohols in heterologous systems. Finally, although the terminal oxidation step producing the aldehyde pheromone component was not identified in this study, accumulation of stable fatty acid and alcohol intermediates may in fact be advantageous for plant-based production systems, as volatile aldehydes can exhibit increased reactivity and potential phytotoxicity in planta. Separation of precursor biosynthesis and final aldehyde conversion may therefore simplify scalable bioproduction workflows.

In summary, we have elucidated the biosynthetic pathway of the sex pheromone of *C. medinalis* and identified the functional desaturase, elongase and reductase. We used this knowledge to guide the reconstruction of the fatty acid metabolic flux of Camelina as a biofactory to produce the fatty acid precursor of the *C. medinalis* major pheromone component. Geographical variation in *C. medinalis* pheromone composition (Kawazu et al., 2000; Rao et al., 1995) further highlights the flexibility of the engineered Camelina platform. Indeed, the simultaneous accumulation of Z13-18, Z11-18, and Z11-16 intermediates provides precursor molecules corresponding to pheromone blends used by different regional populations and may also support production of related pheromone systems, including those of other rice pests such as *Chilo suppressalis*, which uses Z11-16 and Z13-18 as major and minor pheromone components, respectively. Together, our results establish a modular framework for engineering long-chain pheromone biosynthesis in oilseed crops and support the further development of sustainable, plant-based pheromone production systems for integrated pest management.

## 4. Methods

### 4.1 Insect rearing and collection of sex pheromone glands

The colony of *Cnaphalocrocis medinalis* was initially obtained from a paddy field in Huzhou (Zhejiang, China). The insects were maintained in a climate chamber at 27±1 °C and 55±10% relative humidity, and a light:dark photoperiod of 14:10h. For experiments, males and females were kept separately after the pupal stage. Newly emerged adults were held in single-sex cohorts with fresh honey water as food source.

### 4.2 Gas chromatography/ mass spectrometry (GC/MS) analysis of sex pheromone components and fatty acid precursors

For pheromone analyses, the pheromone glands (PG) of 36 to 72h old virgin females were dissected and extracted using 50 μL of heptane (Merck) per 10 pooled glands. Following a 30 min incubation period, the solvent was retrieved and transferred to a new vial for pheromone analysis via GC/MS.

For precursor analysis, the same pooled gland sample was reextracted with a chloroform:methanol (2:1 v/v) (Merck) solution (total lipid extraction) overnight at 25 °C. The extract was then transferred to a new vial, and the solvent was evaporated under a gentle N_2_ flow. Subsequently, the sample was subjected to base methanolysis to transform lipids into fatty acid methyl esters (FAMEs) which were then analyzed by GC/MS.

Pheromone and FAME PG extracts were analysed using an Agilent 5977C mass detector (EI, 70 eV) coupled to an Agilent 8890 series gas chromatograph, fitted with an HP-INNOWax column (30 m × 0.25 mm i.d., 0.25 μm film thickness; Agilent, Technologies, USA). Helium was used as the carrier gas. The GC inlet was set to 250 °C and the oven program to 80 °C for 1 min, increase of 10 °C min^−1^ to 230 °C, held for 10 min. The transfer line temperature was 280 °C. Full scan spectra were recorded in mass range *m/z* 30-400. Selected ion monitoring was performed as described below in 4.3. Data were analysed by the ChemStation software (Agilent, Technologies, USA). Compounds were identified by comparing their retention times and mass spectra to those of synthetic standards.

### 4.3 In vivo labeling

Deuterium-labelled [16,16,16-^2^H_3_] hexadecanoic acid (D_3_-16:Acid), [16,16,16-^2^H_3_] (*Z*)-11-hexadecanoic acid (D_3_-Z11-16:Acid), [18,18,18-^2^H_3_] octadecenoic acid (D_3_-18:Acid) were purchased from Larodan Fine Chemicals, Sweden.

The D_3_-16:acid, D_3_-Z11-16:Acid, and D_3_-18:acid were dissolved individually in dimethylsulphoxide (DMSO) at a concentration of 40 µg µL^-1^. Two-day old insects were anesthetized and 0.2 µL of labelled compound solution was topically applied to the extruded female PG one to 2 hrs into the scotophase. 0.5 h before the pheromone gland extraction. Pheromone gland extraction was performed as described above.

Deuterium incorporation into the pheromone precursor and products was monitored by selected ion monitoring (SIM). Native 16:Me and D_3_-16:Me were monitored using the molecular ion [M]^+^ at *m/z* 270 and 273, respectively. Native Z11-16:Me and D_3_-Z11-16 were monitored using ions at *m/z* 236/268 and *m/z* 239/271, respectively, corresponding to the [M-32]^+^ fragment and molecular ion. Native 18:Me and D_3_-18:Me were monitored using the molecular ion at *m/z* 298 and 301, respectively. Native Z13-18:Me and D_3_-Z13-18:Me were monitored using ions *m/z* 264/296 and *m/z* 267/299, respectively, corresponding to the [M-32]^+^ fragment and molecular ion. Finally, Z13-18:Ald and D_3_-Z13-18:Ald were monitored using at *m/z* 248 and 251, corresponding to the [M-18]^+^ fragment.

### 4.4 Transcriptome sequencing and analysis

Female pheromone gland and abdomen cuticle samples were collected at 36 h post-eclosion. Total RNA was extracted using Trizol Reagent (Thermo Fisher Scientific, United States) following the manufacturer’s protocol. RNA-seq was performed by the Novogene company. Three biological replicates were used for each treatment. Adaptor sequences and low-quality reads were removed using FASTP (Chen et al., 2018). Clean reads were mapped to the *C. medinalis* reference genome (InsectBase (GCA_014851415.1)) using HISAT2 (Kim et al., 2015). The read counts were obtained and normalized by STRINGTIE (Pertea et al., 2015). Principal component analysis (PCA) was performed using the R package VEGAN (Dixon, 2003). Differentially expressed genes (DEGs) were analyzed by the R package EDGER (v.4.22.5) (Robinson et al., 2010).

### 4.5 RNA Extraction and Quantitative RT-qPCR Analysis

For insect samples, total RNA was extracted using Trizol Reagent (Thermo Fisher Scientific, United States) following the manufacturer’s protocol. One microgram of total RNA for each sample was reverse transcribed using the HiScript II Q RT SuperMix (Vazyme). qRT-PCR was performed on the CFX96 Touch (BioRad) using ChamQ SYBR qPCR Master mix (Vazyme). Five independent biological samples were collected and analyzed. Primers used for qRT-PCR are listed in SI Appendix, Table S1. The *C. medinalis* actin mRNA was used as the internal control.

### 4.6 Phylogenetic analysis

Protein sequences were aligned with CLUSTALW, and the tree was constructed using the maximum likelihood method in MEGA5 (Tamura et al., 2011). Bootstrap values (%) obtained from 100 replicates were shown. Sequence abbreviations correspond to species names as follows: Aips, *Agrotis ipsilon*; Aseg*, Agrotis segetum*; Mbra, *Mamestra brassicae*; Hass, *Helicoverpa assulta*; Sexi, *Spodoptera exigua*; Slit, *Spodoptera litura*; *Spodoptera littoralis*; Tni, *Trichoplusia ni*; Lbot, *Lobesia botrana*; Dpun, *Dendrolimus punctatus*; Pxyl, *Plutella xylostella*; Aper, *Antheraea pernyi*; Msex, *Manduca sexta*; Bmor, *Bombyx mori*; Sins, *Streltzoviella insularis*; Tpit, *Thaumetopoea pityocampa*; Pgos, *Pectinophora gossypiella*; Epos, *Epiphyas postvittana*; Avel, *Argyrotaenia velutinana*; Cpar, *Choristoneura parallela*; Cros, *Choristoneura rosaceana*; Csup, *Chilo suppressalis*; Ofur, *Ostrinia furnacalis*; Onub, *Ostrinia nubilalis*; Cpom, *Cydia pomonella*; Lcap, *Lampronia capitella*; Ecau, *Ephestia cautella*; Atra, *Amyelois transitella*; Obru, *Operophtera brumata*; Cpar, *Choristoneura parallela*; Epos, *Epiphyas postvittana*; Sls, *Spodoptera littoralis*; Cher, *Ctenopseustis herana*; Cobl, *Ctenopseustis obliquana*; Dpl, *Danaus plexippus*; Dsac, *Diatraea saccharalis*; Har, *Helicoverpa armigera*; Her, *Heliconius erato*; Hmel, *Heliconius melpomene*; Hvi, *Heliothis virescens*; Opa, O*strinia palustralis;* Osc, ostrinia scapulalis; Oze, *Ostrinia zealis*; Yev, *Yponomeuta evonymella*; Ypa, *Yponomeuta. padella*; Yro, *Yponomeuta. rorrella*.

### 4.7 Plasmid constructs for transient expression

All gene sequences were derived from transcriptome sequencing data and *C. medinalis* reference genome. They were synthesized (Tsingke Biological Technology) and cloned into the plant expression vector pXZP393 by Gateway recombination cloning technology (Invitrogen). After confirming the integrity of the constructs by sequencing, all constructs were transformed into GV3101 *Agrobacterium*. Leaves of 4-wk-old *N. benthamiana* were infiltrated with agrobacterial cells containing the indicated constructs. Detailed procedures were performed as described (Ding et al., 2014).

### 4.8 Fatty acid analysis of *N. benthamiana* leaves

Total lipids were extracted from ca. 0.5 g of leaf tissue using 3.75 mL using methanol/chloroform (2:1, v/v) with internal standard methyl (*Z*)-10-heptadecenoate (Z10-17:Me, 10 µg spiked in), and then acetic acid (0.075 M in water, v/v) was added to produce a biphasic mixture. Tubes were vortexed vigorously and centrifuged. Then, the chloroform phase containing total lipids was transferred to a new glass tube. The solvent was evaporated under nitrogen flow. Then sulphuric acid (2% in methanol, v/v) was added, and the reactant was vortexed vigorously followed by 1 h incubation at 90 °C. After incubation, acetic acid (0.075 M in water, v/v) was added, the mixture was vortexed and heptane was added to extract the FAMEs. FAMEs were analyzed by GC-MS as described above.

### 4.9 Fatty alcohol (OH) analysis of *N. benthamiana* leaves

Total lipids were extracted from ca. 0.5 g of leaf tissue using 3.75 mL of methanol/chloroform (2:1, v/v) with internal standard heptadecanol (17:OH, 10 nmol spiked in), in a glass tissue grinder. The crude extract was transferred to a glass tube and 2 mL of acetic acid (0.075 M in water, v/v) was added to the tube producing a biphasic mixture. Tubes were vortexed vigorously and centrifuged at 2000 rpm for 2 min. The ca. 1 mL of chloroform phase containing the total lipids was transferred to a new glass tube, which was then concentrated to ca. 50 mL under gentle nitrogen flow.

The concentrated sample was subsequently loaded onto a TLC plate (Silica gel 60, Merck, Germany) and separated with a mobile phase of hexane/diethyl ether/acetic acid (85:15:1, v/v/v). The bands were visualized by spraying water, and target gel areas were collected separately into 4 mL vials and extracted with 1 mL of methanol/chloroform (2:1, v/v) in sonication bath for 30 s. The extract was then centrifuged at 2000 rpm for 2 min. The supernatant was transferred to a new tube and 1 mL acetic acid (0.075 M in water, v/v) was added to partition the lipids into chloroform. The chloroform phase containing alcohols, was transferred into new tubes for subsequent GC/MS analysis as described above.

### 4.10 Generation and characterization of transgenic Camelina plants

Camelina (*Camelina sativa*) var. *Sunesson* was used as the transformation background. Plants were grown in a growth chamber under 14-h light (25 °C) and 10-h dark (20 °C) photoperiod, with a light intensity of ∼ 140 µmol m^−2^ s^−1^ (LED light) and 60% relative humidity. The Cuphea *FatB1* thioesterase (*FatB1*) gene (KC675176.1), *CsupYPAQ* gene (MN453822.1), *CsaKCS1* gene (Csa03g002110.1) and Red Fluorescent Protein from Discosoma (*DsRED*) (UXW73585.1) were introduced into the binary vector pBinGlyBar using Gateway cloning. *FatB1* was placed under control of glycinin promoter. *CsupYPAQ* and *CsaKCS1* were controlled by napin promoter. *DsRED* was controlled by Cauliflower Mosaic Virus 35S promoter. *Csa_Z11-16* transgenic line contains *FatB1 and CsupYPAQ*. *Csa_Z13-18* transgenic line contains *FatB1*, *CsupYPAQ*, *CsaKCS1* and *DsRED*. The pBinGlyBar binary vector contains a constitutively expressed resistance gene for the herbicide phosphinothricin (or Basta). After confirming the integrity of the constructs by sequencing, the resulting binary vector was transformed into GV3101 *Agrobacterium*. *A. tumefaciens* cells harbouring the binary vector were transformed by floral infiltration into the *Camelina* as previously described (Lu and Kang, 2008). Positive lines from collected T1 seeds were screened for Basta resistance and Red Fluorescence. T2 seeds from these lines were screened by GC/MS.

FAMEs were generated by grinding pools of 10 seeds per plant (N=8) in 1 mL 2% sulphuric acid in methanol followed by incubation for 1 h at 90 °C in a 4 mL glass vial. After cooling, 1 mL water was added, the mixture was vortexed and then 1 mL heptane was added to extract the FAMEs. Finally, the heptane phase containing the FAMEs was transferred to1.5 mL autosampler vials for GC/MS analysis as described above.

## Author Contributions

ML, BD and CL, conceived the study. ML and CL obtained funding. HL performed the *C. medinalis* capture and rearing. HL and ML conducted the insect sample collection and in vivo labeling experiments. ML carried out the transcriptome and gene expression analysis. ML, BD and YX carried out vector design and construction. ML and HL performed the *N. benthamiana* transient expression experiment. PH, CT, HJ and ML performed floral dip transformation, plant cultivation, sample analysis and greenhouse propagation in Hangzhou. ML., CL, JML. and HL analyzed the data. JML, HL and ML completed the design and standardization of figures. ML, BD and YX drafted the manuscript. CL, JML and HW edited the manuscript. All authors approved the final version.

## Acknowledgements

We thank Yaobin Lu from Xianghu Laboratory, Yiwei Zhang from ZJU-Hangzhou Global Scientific and Technological Innovation Center for technical assistance, Xi Li, Chengqi Zhu from Xianghu Laboratory, and Xiaomu Qiao from Zhejiang A&F University for valuable discussion. Figdraw for drawing assistance. This work was supported by the National Natural Science Foundation of China (grant no. 32402446), the Key R&D Program of Zhejiang (grant no. 2024SSYS0102), the Hangzhou Natural Science Foundation Project (No. 2024SZRYBC40001).

## Data Availability Statement

The RNA-Seq data reported in this paper have been deposited in the Genome Sequence Archive at the BIG DataCenter (http://bigd.big.ac.cn/gsa), Beijing Institute of Genomics (BIG), Chinese Academy of Sciences, under accession nos. CRA039891.

## Supporting Information

Additional supporting information can be found online in the Supporting Information section. **Figure S1:** Fatty acid profile of female *C. medinalis* pheromone gland. **Figure S2:** Tissue-specific expression profiles of candidate pheromone biosynthetic genes in female *C. medinalis*. **Figure S3:** Phylogenetic analysis of Lepidoptera fatty acyl desaturases (FADs). **Figure S4:** Phylogenetic analysis of plant fatty acid elongases (KCS family). **Figure S5:** Functional screening of plant elongases in *Nicotiana benthamiana*. **Figure S6:** Growth and yield-related traits of transgenic *Camelina sativa* lines. **Table S1:** Primers sequences used in this study.

