## Supplementary Data for "Paving the way for plant-based bioproduction of (Z)-13-octadecenoic moth pheromone compounds"

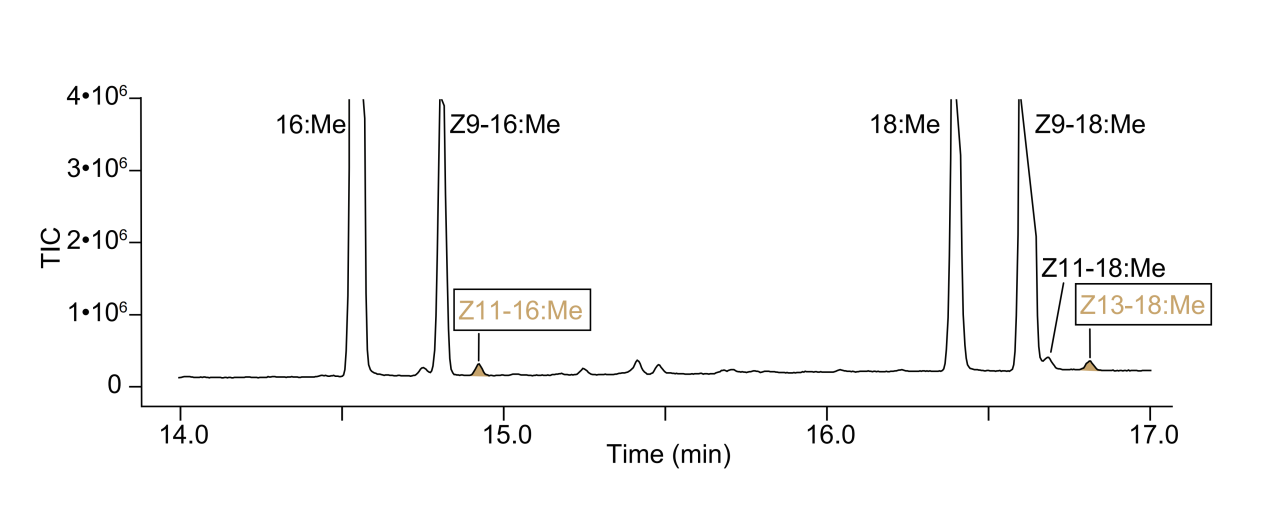

**Figure S1.** **Fatty acid profile of female *C. medinalis* pheromone gland**.

Total ion chromatogram (TIC) of fatty acid methyl esters (FAMEs) derived from base-methanolized pheromone glands of *C. medinalis* females. In addition to common fatty acid derivatives, compounds associated with pheromone components are highlighted. Compound identities were confirmed by comparison of retention time and mass spectra of synthetic standards. Abbreviations: 16:Me: methyl hexadecanoate: Z9-16:Me: methyl (*Z*)-9-hexadecenoate; Z11-16:Me: methyl (*Z*)-11-hexadecenoate; 18:Me: methyl-octadecenoate; Z9-18:Me: methyl (*Z*)-9-octadecenoate; Z11-18:Me: methyl (*Z*)-11-octadecenoate; Z13-18:Me: methyl (*Z*)-13-octadecenoate.

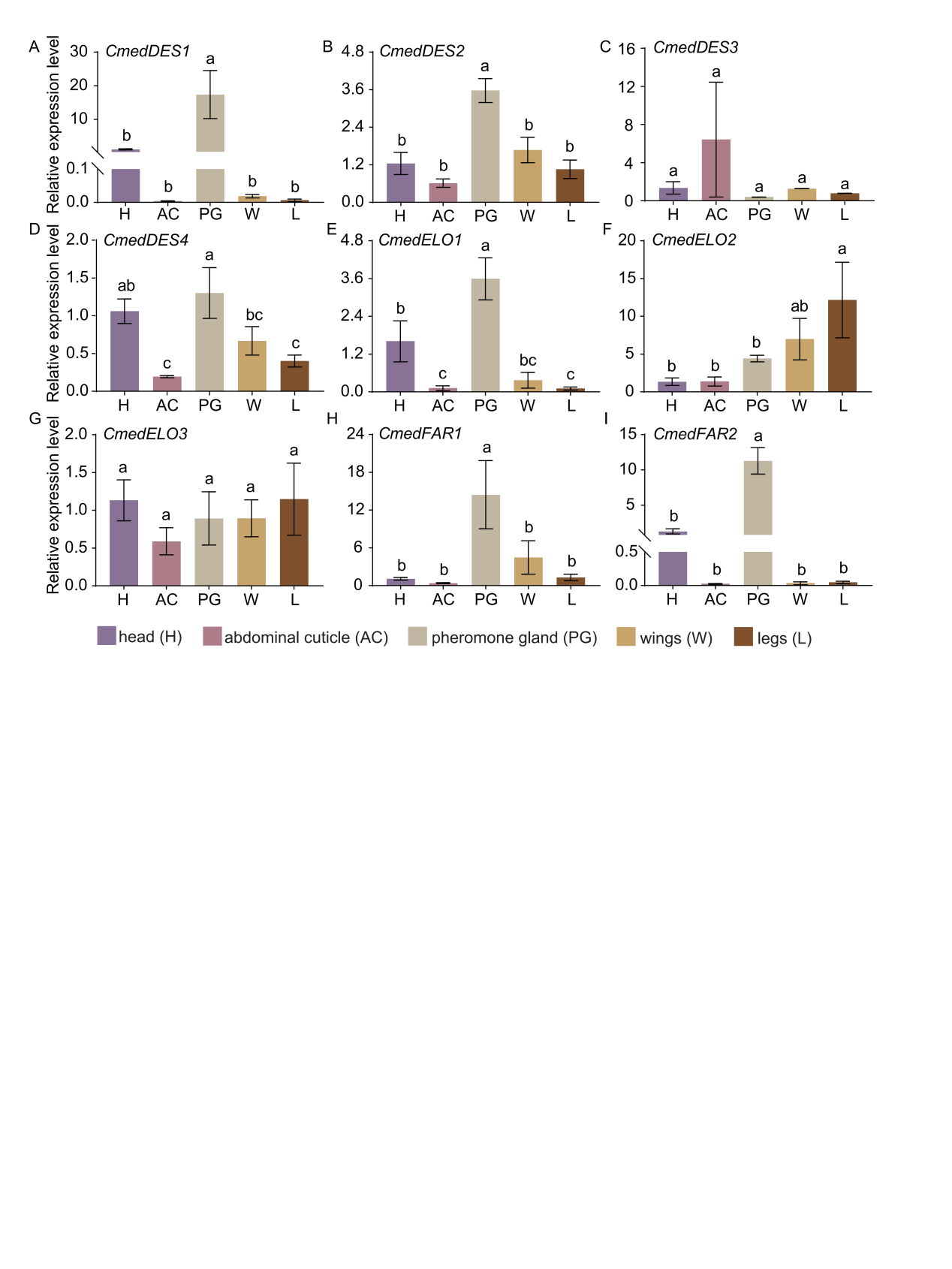

**Figure S2. Tissue-specific expression profiles of candidate pheromone biosynthetic genes in female *C. medinalis*.**

(A–I) Relative expression levels of CmedDES1 (A), CmedDES2 (B), CmedDES3 (C), CmedDES4 (D), CmedELO1 (E), CmedELO2 (F), CmedELO3 (G), CmedFAR1 (H), and CmedFAR2 (I) in different female tissues. Data represent mean values ± SE (n = 4–5). H, head; AC, abdominal cuticle; PG, pheromone gland; W, wing; L, leg. Different letters indicate statistically significant differences among tissues (one-way ANOVA followed by Tukey’s multiple comparison test; P < 0.05).

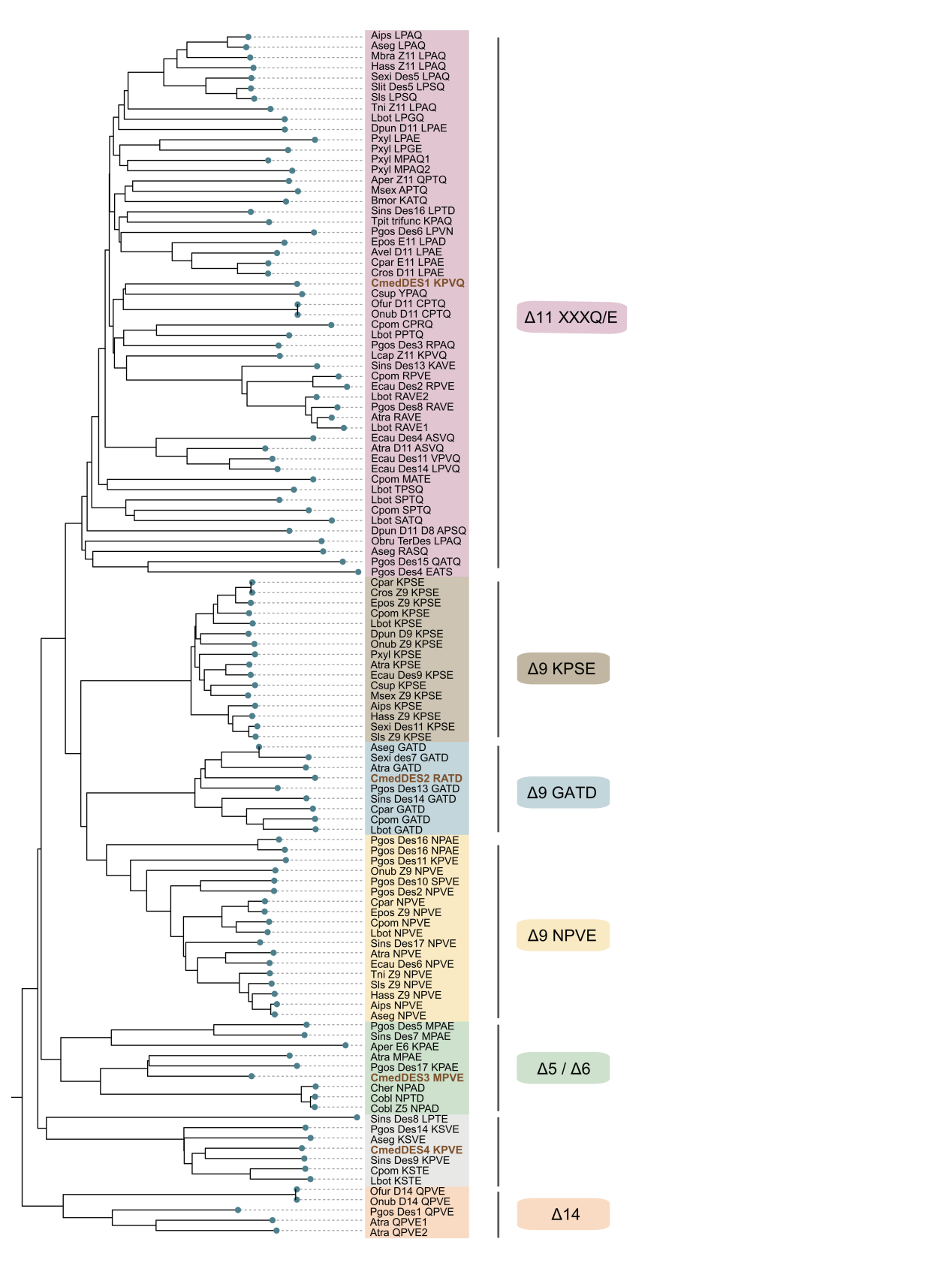

**Figure S3. Phylogenetic analysis of Lepidoptera fatty acyl desaturases (FADs).**

Maximum-likelihood phylogenetic tree of fatty acyl desaturase (FAD) proteins from *Cnaphalocrocis medinalis* and other Lepidoptera species. CmedDES sequences are highlighted in purple, whereas homologous sequences from other species are shown in black. Putative functional classes are indicated by colored labels corresponding to characteristic biochemical activities and consensus signature motifs (e.g., KPSE, NPVE, GATD). Major groups, including Δ9, Δ11/Δ10 bifunctional, Δ5/Δ6, and Δ14 desaturases, are indicated. Sequence abbreviations correspond to species names as follows: Aips, *Agrotis ipsilon*; Aseg*,* *Agrotis segetum*; Mbra, *Mamestra brassicae*; Hass, *Helicoverpa assulta*; Sexi, *Spodoptera exigua*; Slit, *Spodoptera litura*; *Spodoptera littoralis*; Tni, *Trichoplusia ni*; Lbot, *Lobesia botrana*; Dpun, *Dendrolimus punctatus*; Pxyl, *Plutella xylostella*; Aper, *Antheraea pernyi*; Msex, *Manduca sexta*; Bmor, *Bombyx mori*; Sins, *Streltzoviella insularis* ; Tpit, *Thaumetopoea pityocampa*; Pgos, *Pectinophora gossypiella*; Epos, *Epiphyas postvittana*; Avel, *Argyrotaenia velutinana*; Cpar, *Choristoneura parallela*; Cros, *Choristoneura rosaceana*; Csup, *Chilo suppressalis*; Ofur, *Ostrinia furnacalis*; Onub, *Ostrinia nubilalis*; Cpom, *Cydia pomonella*; Lcap, *Lampronia capitella*; Ecau, *Ephestia cautella*; Atra, *Amyelois transitella*; Obru, *Operophtera brumata*; Cpar, *Choristoneura parallela*; Epos, *Epiphyas postvittana*; Sls, *Spodoptera littoralis*; Cher, *Ctenopseustis herana*; Cobl, *Ctenopseustis obliquana*;

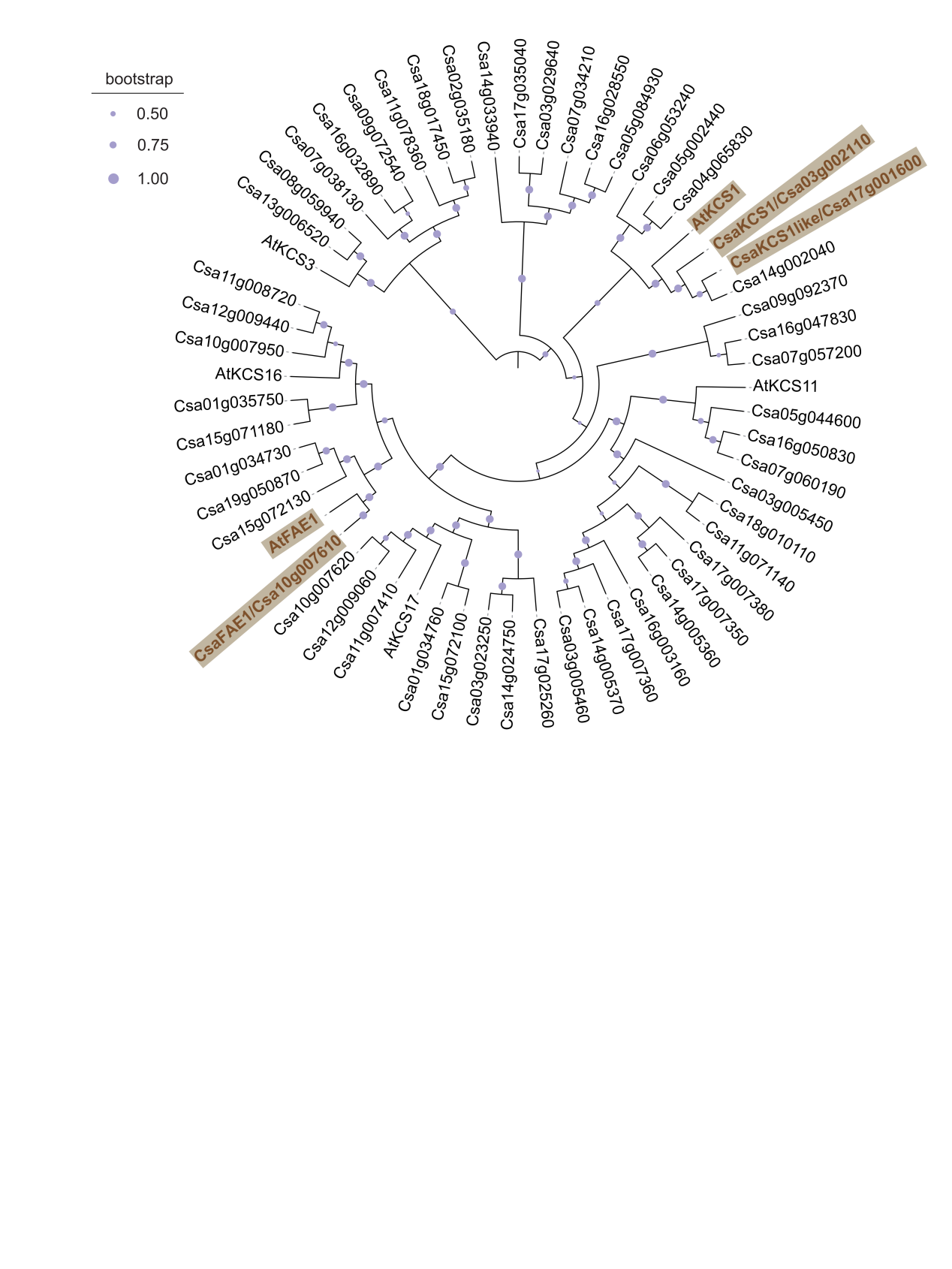

**Figure S4. Phylogenetic analysis of plant fatty acid elongases (KCS family).**

Maximum-likelihood phylogenetic tree of 3-ketoacyl-CoA synthases (KCS) from *Camelina sativa* and *Arabidopsis thaliana*. Candidate plant elongases selected for functional characterization are highlighted in purple, whereas other sequences are shown in black. Known KCS/FAE proteins, namely AtKCS1 (AT1G01120) and AtFAE1 (AtKCS18; AT4G34520) are included as references for functional classification.

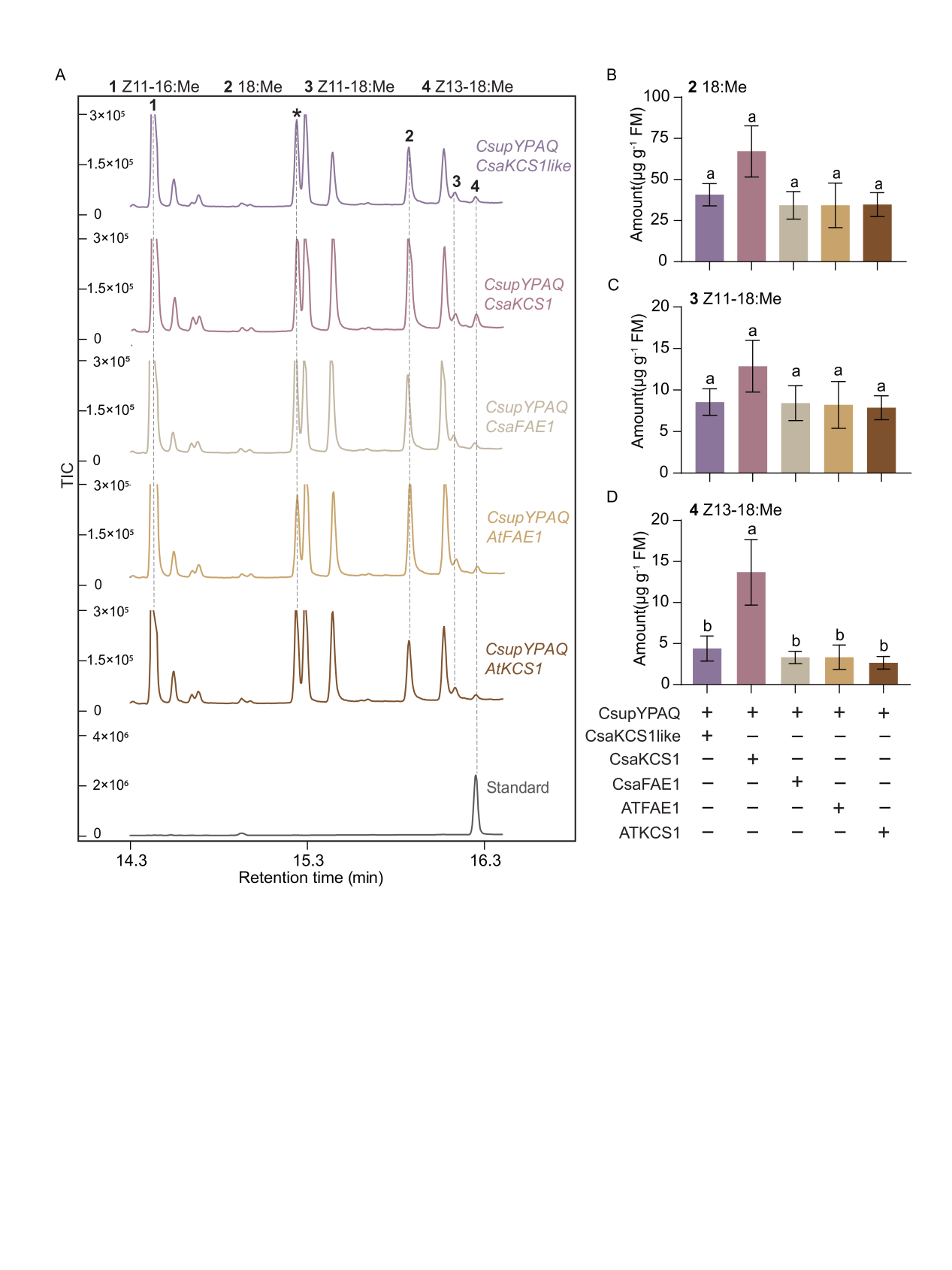

**Figure S5. Functional screening of plant elongases in *Nicotiana benthamiana*.**

(A) GC–MS analysis of fatty acid methyl ester (FAME) profiles from *N. benthamiana* leaves co-expressing CsupYPAQ with plant elongases: CsaKCS1-like, CsaKCS1, CsaFAE1, AtFAE1, and AtKCS1. Total ion chromatograms (TICs) are shown for each combination. Peaks are annotated as follows: Z11-16:Me (1), 18:Me (2), Z11-18:Me (3), and Z13-18:Me (4). The asterisk indicates the internal standard (Z10-17:Me, 10 μg spiked in). TIC of an authentic Z13-18:Me standard. (B–D) Quantification of FAME products in *N. benthamiana* leaves expressing the indicated constructs. Mean levels (± SE, n = 5) of 18:Me (B), Z11-18:Me (C), and Z13-18:Me (D) are shown. Metabolite levels were measured 72 h after agroinfiltration. Different letters indicate statistically significant differences among treatments (one-way ANOVA followed by Tukey’s multiple comparison test; P < 0.05).

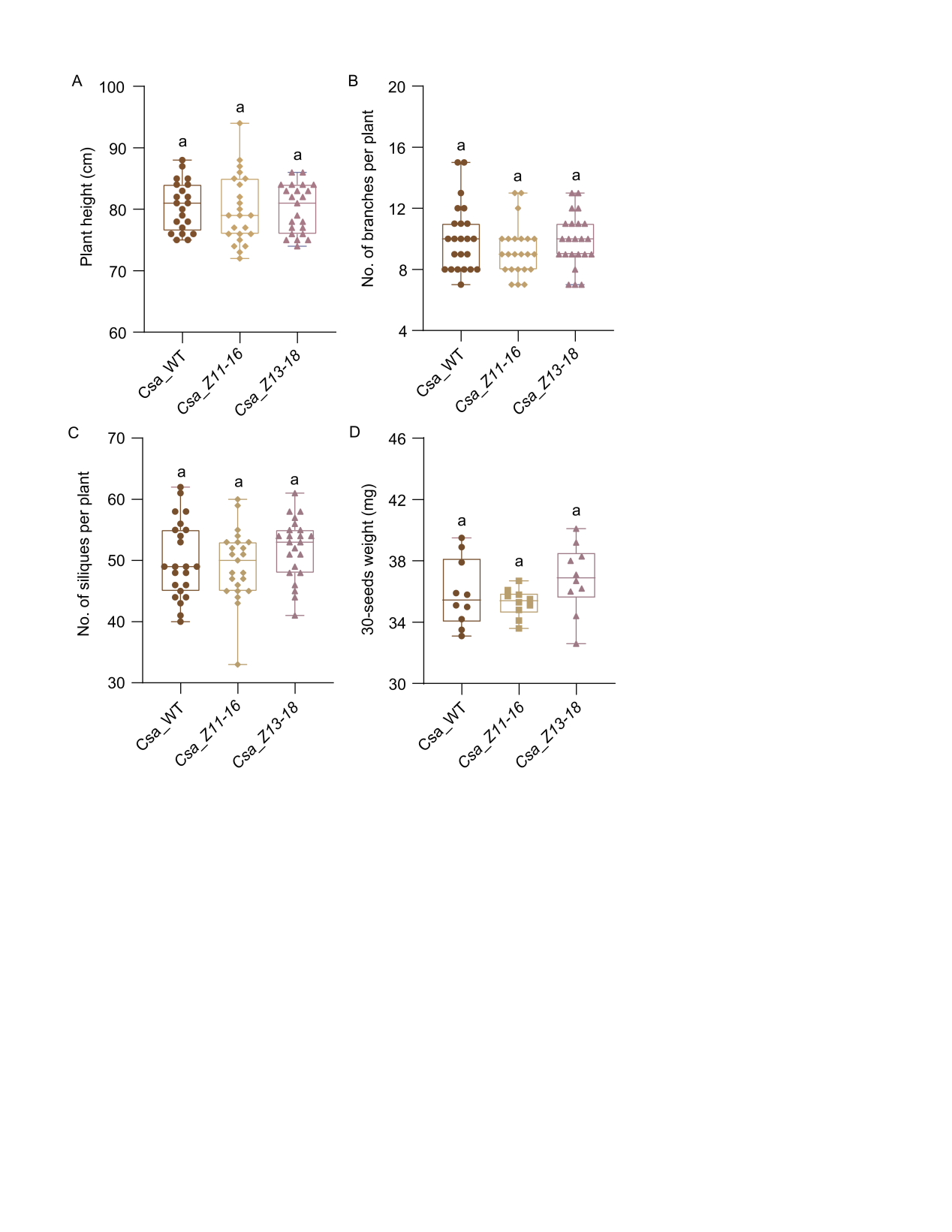

**Figure S6. Growth and yield-related traits of transgenic *Camelina sativa* lines.**

(A–D) Phenotypic analysis of wild-type (WT), Csa_Z11-16, and Csa_Z13-18 transgenic *Camelina sativa* plants. Mean plant height (A), number of branches per plant (B), number of siliques on the main branches per plant (C), and 30-seeds weight (D) are shown. Data represent mean values ± SE (n = 21). Plants were analyzed at 80 days after growth under standard conditions. Different letters indicate statistically significant differences among genotypes (one-way ANOVA followed by Tukey’s multiple comparison test; P < 0.05).

Table S1 Primer used in this study

| Gene ID | Primer | Sequence (5’-3’) | Purpose |
| --- | --- | --- | --- |
| Cmed070400 | CmedDES1-RT-F | TTGGCGTTCCAAAATACGGC | RT-qPCR |
|  | CmedDES1-RT-R | GGAACCGCAAAACAGGGTTG | RT-qPCR |
| Cmed116590 | CmedDES2-RT-F | TGGCCTTCCAGAACCACATC | RT-qPCR |
|  | CmedDES2-RT-R | GTCGATGGTGGACCCCTTTT | RT-qPCR |
| Cmed116640 | CmedDES3-RT-F | ACCGGGTCCATCACAAGTTC | RT-qPCR |
|  | CmedDES3-RT-R | GATGATGAAGCAGCACAGCG | RT-qPCR |
| Cmed116600 | CmedDES4-RT-F | TGTGGATACCGTGGCACTTC | RT-qPCR |
|  | CmedDES4-RT-R | CGCAAAACTGTTGACGCAGA | RT-qPCR |
| Cmed153350 | CmedELO1-RT-F | GTCGAACTAGCTGATCCCCG | RT-qPCR |
|  | CmedELO1-RT-R | CCATCCTGCATCCAAACCCT | RT-qPCR |
| Cmed153310 | CmedELO2-RT-F | TGGAAGAAGCACCTCACCAC | RT-qPCR |
|  | CmedELO2-RT-R | GATAGCCGCAGTCGTAGACC | RT-qPCR |
| Cmed153380 | CmedELO3-RT-F | TACAATGCCGTCCAAGTGCT | RT-qPCR |
|  | CmedELO3-RT-R | GCTCGTTGTTCAGCAAGCAA | RT-qPCR |
| Cmed114440 | CmedFAR1-RT-F | ACTGCATTCTCCAACTCGCA | RT-qPCR |
|  | CmedFAR1-RT-R | CTCGGCTAGTGCTTTGGTGA | RT-qPCR |
| Cmed122000 | CmedFAR2-RT-F | GGCCATGCTGTGTCAAAAGG | RT-qPCR |
|  | CmedFAR2-RT-R | CTCGAGGCCAAGCATACGAT | RT-qPCR |
| Cmed118340 | Actin-RT-F | CGAGCGTGGTTACTCATTCA | RT-qPCR |
|  | Actin-RT-R | ATGACTTCTCGAGCGAGCTG | RT-qPCR |
